# Lipid Imbalance Generates Golgi Whorls that Sequester Small GTPases

**DOI:** 10.64898/2026.07.23.740422

**Authors:** Qiuxin Zhang, Weiyi Tan, Jiaqi Guo, Isabela Ashton-Rickardt, Joshua Harrison, Susan J. Birren, Jun Liu, Jie Zhou, Jennifer Lippincott-Schwartz, Bing Xu

## Abstract

Small GTPases are generally viewed as organizers of organelle identity, with different families recruited to specific membranes by compartment-selective targeting mechanisms. Here, we identify a membrane state that reverses this relationship. Excessive peptide S-palmitoylation at Golgi generates multilamellar, filipin-poor whorls that recruit ARF, Rab, and Rho family GTPases normally associated with distinct cellular compartments. The whorls simultaneously excluded Golgi transmembrane residents, coat proteins, ER proteins, a GPI-anchored protein, and other palmitoylated proteins, demonstrating that they were selective membrane compartments rather than nonspecific protein aggregates. Preventing myristoylation of ARF6 or geranylgeranylation of Rab11a strongly reduced recruitment, whereas prenylation alone was insufficient, indicating recognition of a composite membrane-targeting code. Lowering cellular cholesterol promoted whorl formation, whereas cholesterol supplementation suppressed it. Chemically distinct lipid perturbations generated GTPase-positive Golgi whorls, suggesting convergence on a shared membrane-remodeling state. As whorls accumulated, endocytic recycling and anterograde trafficking declined, survival signaling decreased, and viability became poorly reversible. These findings show that lipid imbalance can redirect small GTPases across normal organelle boundaries, converting Golgi-derived membranes into selective sinks for trafficking regulators.

## Introduction

Small GTPases are central determinants of organelle identity. ARF, Rab, and Rho proteins occupy different membrane compartments, where their spatially restricted activation coordinates vesicle formation, fusion, cytoskeletal organization, and signaling (*1*). This framework assumes that compartment-specific regulatory machinery and lipid cues maintain the separation of these GTPase systems. Whether a membrane can enter a physical state that overrides this separation and recruits multiple GTPase families to the same ectopic compartment is unknown.

S-palmitoylation is generally understood as one of the mechanisms that preserves this spatial order. The reversible attachment of a 16-carbon fatty acid to cysteine residues (*2*) regulates protein membrane association, trafficking, stability, and signaling (*3, 4*), allowing proteins to cycle between membrane compartments with temporal and spatial precision. Dysregulated S-palmitoylation contributes to cancer (*5-7*), neurodegeneration (*8, 9*), and metabolic disease (*2*), and it modulates stress and cell-death pathways death pathways including pyroptosis, ferroptosis, necroptosis, and autophagy-associated responses (*10-16*). These studies establish S-palmitoylation as a powerful determinant of protein localization and cell fate. They leave open a more disruptive possibility: when palmitoylation becomes excessive at one organelle, can the membrane itself acquire a new identity and capture proteins that are normally segregated elsewhere?

The Golgi apparatus provides a stringent setting in which to test this possibility because it is a major hub for S-palmitoylation, lipid sorting, and membrane trafficking. Golgi-localized ZDHHC acyltransferases and accessory proteins such as GOLGA7 concentrate S-palmitoylation activity at this organelle (*4, 17-20*), where palmitoylation promotes anterograde cargo sorting and regulates the recycling of peripheral membrane proteins between the Golgi and other compartments (*4, 21*). Because Golgi combines concentrated acylation activity with continuous membrane influx and efflux, the prevailing expectation is that lipid or palmitoylation imbalance would disrupt cargo flow or Golgi integrity. Consistent with this view, Golgi disruption impairs vesicle transport, disturbs lipid homeostasis, and can precede cell death (*22*), whereas excessive (*16*) or spatially dysregulated (*14*) palmitoylation is linked to cellular stress. What remains unknown is whether these perturbations can produce a distinct Golgi-derived membrane state that overrides native GTPase targeting.

Multilamellar membrane whorls appear, at first, to be an unlikely candidate for such a state. They have been observed mainly in the endoplasmic reticulum (ER) or lysosome system during protein overload (*23*), lipid storage defects (*24*), drug toxicity (*25*), or dysregulated autophagy (*26*), and are commonly interpreted as passive or nonspecific membrane accumulations. Golgi-derived whorls have been reported only rarely (*27, 28*), leaving unresolved whether they represent organelle collapse or compositionally specialized membrane compartments. If whorls are merely membrane debris, they should not selectively recruit signaling proteins according to shared targeting features. We therefore asked whether a localized Golgi lipid imbalance can generate multilamellar structures and redistribute small GTPases across their normal organelle boundaries.

Here, we find that excessive peptide S-palmitoylation does not simply disorganize the Golgi. Instead, it separates Golgi-derived membranes into multilamellar, cholesterol-poor whorls and cholesterol-retaining puncta. The whorls selectively recruit members of the ARF, Rab, and Rho families while excluding canonical Golgi and ER residents, coat proteins, a GPI-anchored protein, and other palmitoylated proteins. Mutational and pharmacological experiments show that intact GTPase lipid anchors are required for efficient recruitment. Lowering cellular cholesterol promotes whorl formation, whereas cholesterol supplementation suppresses it, linking whorl biogenesis to local sterol availability.

Chemically distinct lipid-imbalance agents, including CIL56 (*14*) and tegavivint (*29*), and 7-ketocholesterol (7-KC) (*30, 31*), produce related ARF6-positive whorls, providing independent convergence on a recurring membrane-remodeling response rather than a peptide-specific artifact. As whorls accumulate, endocytic recycling and anterograde transport decline, cytoskeletal and lipid organization change, AKT–FOXO3a signaling is suppressed, and cell viability decreases through a pathway that resists inhibitors of several canonical death programs. Together, these findings reveal a limit to the compartment-specific view of small-GTPase localization: under severe lipid imbalance, membrane composition can override native organelle targeting and convert Golgi-derived membranes into selective sinks for trafficking regulators.

## Results

### Golgi-localized peptide S-palmitoylation converts Golgi membranes into multilamellar membrane whorls

To create a localized excessive S-palmitoylation at the Golgi, we used the self-assembling peptide precursor AcS-bb-NBD (**1**, Fig. 1A). Its acetyl thioester undergoes intracellular deacetylation (*16, 32*), which exposes a free thiol; cellular acyltransferases (*4*) then S-palmitoylate the thiol to generate palmS-bb-NBD. LC-HRMS analysis of lysates from HeLa cells treated with **1** detected the ionic and isotopic pattern predicted for the palmitoylated product (Fig. 1B). At 0.5 μM, the NBD signal concentrated at the Golgi and overlapped with GalT and Giantin (Fig. 1C). These data establish **1** as a tool for increasing the abundance of palmitoylated peptide at Golgi membranes.

**Figure 1.**
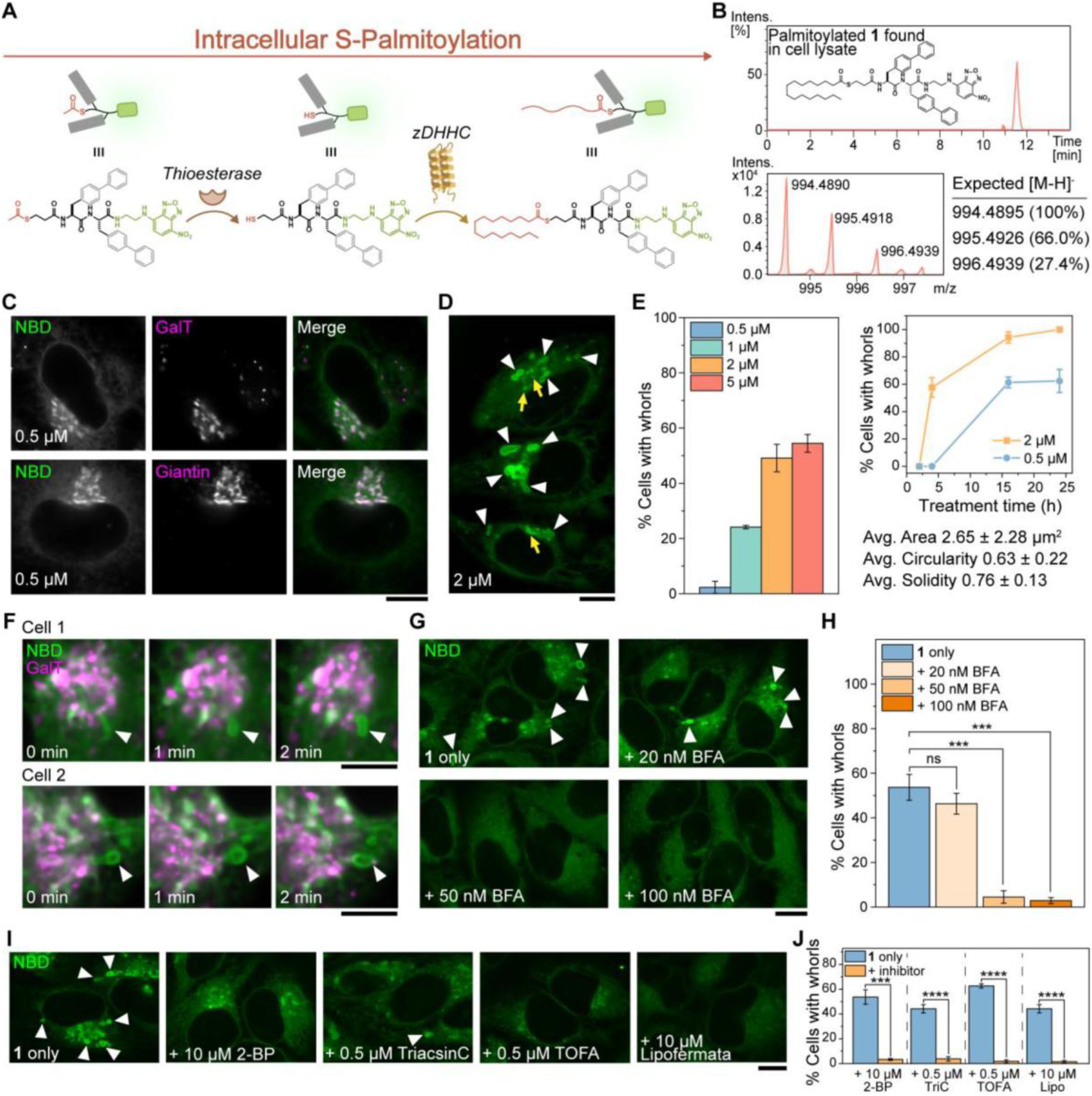
Golgi-localized peptide S-palmitoylation converts Golgi membranes into multilamellar whorls. (**A**) Schematic of intracellular S-palmitoylation of the peptide precursor AcS-bb-NBD (**1**). Cellular thioesterases deacetylate the thioester to expose a free thiol, which is then S-palmitoylated by Golgi-resident ZDHHC acyltransferases to yield palmS-bb-NBD. (**B**) Extracted ion chromatogram (top) and high-resolution mass spectrum (bottom) from LC-HRMS analysis of HeLa cell lysates treated with **1**, confirming in situ formation of the palmitoylated species. (**C**) Representative CLSM images of HeLa cells transiently transfected with mCh-GalT (top) or mScarlet-Giantin (bottom) and treated with **1** (0.5 μM, 4 h), showing colocalization of NBD signal with the trans-Golgi marker GalT or the cis/medial-Golgi marker Giantin. (Scale bar = 5 μm) (**D**) Representative CLSM image of HeLa cells treated with **1** (2 μM, 4 h). White arrowheads indicate multilamellar whorl-like structures; yellow arrows indicate NBD-positive puncta. (Scale bar = 10 μm) (**E**) Quantification of whorl formation in HeLa cells treated with **1**. Left: percentage of cells containing whorls across concentrations (4 h treatment). Right: percentage of cells containing whorls over time at 0.5 μM and 2 μM. Whorl morphology parameters (mean ± s.d., n = 399): area 2.65 ± 2.28 μm^2^, circularity 0.63 ± 0.22, solidity 0.76 ± 0.13. (**F**) Time-lapse CLSM images of two representative HeLa cells transiently transfected with EBFP-GalT (magenta) and treated with **1** (2 μM), showing whorl emergence (white arrowheads) from existing GalT-positive Golgi stacks over a 2-minute window. (Scale bar = 5 μm) (**G**) Representative CLSM images of HeLa cells co-treated with **1** (2 μM, 4 h) and Brefeldin A (BFA) at the indicated concentrations. White arrowheads indicate whorls. (Scale bar = 10 μm) (**H**) Quantification of whorl prevalence in HeLa cells co-treated with **1** (2 μM, 4 h) and BFA at the indicated concentrations. (**I**) Representative CLSM images of HeLa cells pretreated with the indicated lipid-pathway inhibitors for 30 min, then co-treated with **1** (2 μM) plus inhibitor for 4 h. Inhibitors target S-acylation (2-BP, 10 μM), long-chain fatty acid activation (triacsin C, 0.5 μM), acetyl-CoA carboxylase (TOFA, 0.5 μM), and fatty acid uptake (lipofermata, 10 μM). White arrowheads indicate whorls. (Scale bar = 10 μm) (**J**) Quantification of whorl prevalence in HeLa cells treated with **1** alone or **1** plus the indicated inhibitors as in (I). Data are mean ± s.d. from 3 independent experiments, with 4 cells or fields analyzed per condition. Statistical significance was determined by Student’s t test: **P < 0.01, ***P < 0.001, ****P < 0.0001; ns, not significant.

Rather than simply increasing Golgi fluorescence, raising the concentration of **1** produced an unexpected morphological transition. At 2 μM or higher, cells developed irregular ring-like structures 1–5 μm in diameter with NBD-bright rims, dim centers, and nearby NBD-bright puncta (Fig. 1D). Whorl prevalence increased with dose and treatment time (Fig. 1E and Fig. S1). Area, circularity, and solidity followed reproducible distributions, which distinguished the structures from random intracellular aggregates (Fig 1E and Fig. S2). Live-cell imaging of EBFP-GalT-expressing cells captured NBD-positive whorls emerging from or immediately adjacent to GalT-positive Golgi membrane within 2 min (Fig. 1F). Time-lapse imaging further showed transient contact between forming whorls and microtubule and actin filaments (Fig. S3), suggesting that the cytoskeleton may facilitate whorl biogenesis. Similar structures formed in several mammalian cell lines, primary mouse neurons and glia, and in *C. elegans* (Fig. S4), indicating that diverse cellular contexts support this response.

To distinguish active membrane remodeling from intracellular peptide precipitation, we tested whether whorl formation required Golgi integrity and fatty-acid metabolism. Brefeldin A (BFA) (*33*), which disrupts Golgi organization by inhibiting ARF guanine nucleotide exchange factors, suppressed whorl formation in a concentration-dependent manner (Fig. 1G, H). Inhibitors of S-acylation (2-BP), long-chain fatty acid activation (triacsin C), acetyl-CoA carboxylase (TOFA), or fatty acid uptake (lipofermata) also reduced whorl formation during treatment with **1** (Fig. 1I, J), as did the Golgi-disrupting agent golgicide A (Fig. S5). Time-lapse imaging showed that co-treatment with 2-BP, or with the thioesterase inhibitors DC661 or ML211, suppressed the real-time appearance of NBD-positive puncta and whorls over the first 30 min of treatment with **1** (Fig. S6A, B). In contrast, inhibiting ARF6 activation with NAV-2729 or bragsin1 barely reduced whorl formation (Fig. S5), indicating that ARF6 GTPase activity is dispensable for whorl biogenesis under these conditions. Although the effective inhibitors act at different points in lipid metabolism, their concordant effects show that whorl formation depends on an intact Golgi and on pathways that supply and transfer fatty acyl groups, rather than on ARF6 catalytic activity itself.

Once formed, whorls responded differently to the same manipulations. Washing out **1** and replacing the medium with 2-BP, lipofermata, TOFA, or triacsin C reduced whorl abundance in a dose-dependent manner, whereas BFA did not (Fig. S6C, D, G, H; Fig. S7), indicating that whorl clearance depends on continued lipid processing rather than on ARF-GEF activity. By contrast, replacing the medium with the thioesterase inhibitors ML211 or DC661, unexpectedly, increased whorl abundance in a dose-dependent manner (Fig. S6C, E, F), suggesting that continuous palmitoylation cycling normally contributes to whorl turnover. These findings argue against simple peptide precipitation and instead support an active, metabolically coupled membrane-remodeling response to Golgi-localized S-palmitoylation.

### Golgi whorls override organelle identity and selectively recruit ARF-, Rab-, and Rho-family GTPases

ARF-, Rab-, and Rho-family GTPases bind membranes through lipid anchors(*34*) (*35*), amphipathic helices (*36*), and electrostatic interactions (*1*), but compartment-specific cues normally keep these families spatially separated. We therefore asked whether the Golgi whorl state preserved or overrode that separation. We treated HeLa cells expressing fluorescently tagged GTPases with **1** and analyzed their distributions by confocal microscopy. Fluorescent protein alone did not accumulate on whorls (Fig. S8), showing that recruitment required features of the GTPase construct.

Contrary to the expected compartment specificity, compound **1** recruited GTPases from each family to the same Golgi-derived structures. ARF1 and Sar1b accumulated on whorl membrane (Fig. 2A) and ARF3, ARF4, ARF5, and ARF6 also redistributed to whorls (Fig. 2A, D; Fig. S9). This redistribution was especially unexpected for ARF6, which normally functions mainly at the plasma membrane and recycling endosomes. Rab4a, Rab5, Rab7a, Rab8a, Rab9a, and Rab11a likewise accumulated on whorl membranes despite their associations with distinct trafficking compartments (Fig. 2B, D; Figs. S9, S10). The Rho-family GTPases CDC42 and Rac1 also localized to whorls despite their predominant functions at the plasma membrane and actin-rich structures (Fig. 2B). Compound **1** recruited mCh-Rab11a to whorls in T98G, PANC-1, SH-SY5Y, HepG2, H460, and A431 cells (Fig. 2H), extending the response across cell types.

**Figure 2.**
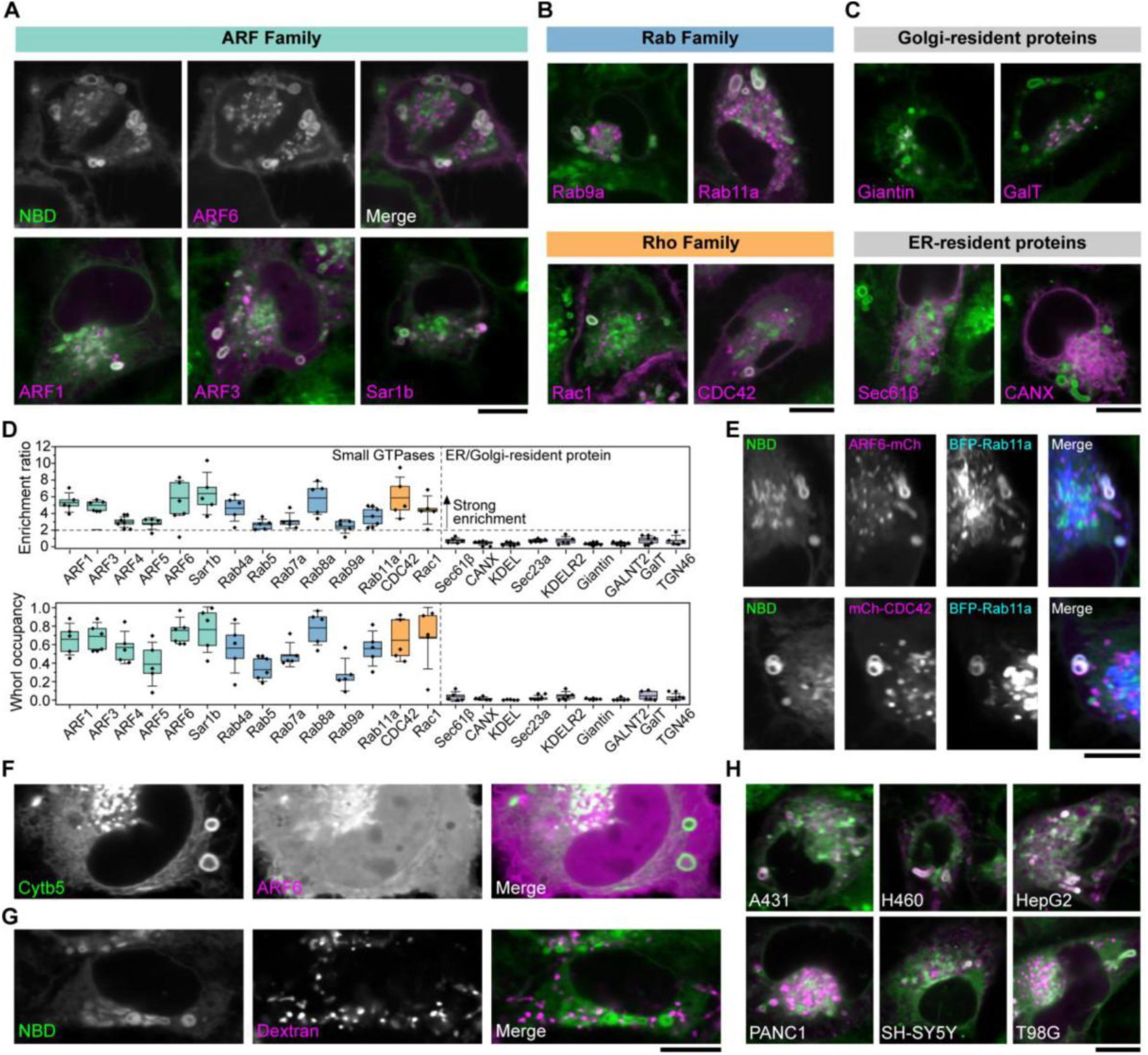
Golgi whorls override organelle identity and selectively sequester ARF, Rab, and Rho-family GTPases. (**A**) Representative CLSM images of HeLa cells transiently transfected with RFP-tagged ARF family GTPases and treated with **1** (1 μM, 4 h). NBD, ARF6, and merge channels are shown for ARF6 to establish the color convention used throughout (green, NBD; magenta, GTPase); merged images only are shown for ARF1, ARF3, and Sar1b. (Scale bar = 10 μm) (**B**) Representative CLSM images (merge) of HeLa cells transiently transfected with RFP-tagged Rab family (Rab9a, Rab11a) or Rho family (Rac1, CDC42) GTPases and treated with **1** (1 μM, 4 h). (Scale bar = 10 μm) (**C**) Representative CLSM images (merge) of HeLa cells transiently transfected with RFP-tagged Golgi-resident (Giantin, GalT) or ER-resident (Sec61β, CANX) proteins and treated with **1** (1 μM, 4 h), showing retention of canonical organelle distribution and absence from whorl membranes. (Scale bar = 10 μm) (**D**) Quantification of whorl recruitment across all proteins tested. Top: enrichment ratio, calculated as the mean fluorescence intensity of the indicated protein at whorls normalized to its mean cytosolic intensity. Bottom: whorl occupancy, calculated as the fraction of whorls per cell positive for the indicated protein. Boxes indicate the interquartile range, center lines indicate the median, and whiskers indicate the 5th and 95th percentiles. n = 5 independent cells per condition. (**E**) Representative CLSM images of HeLa cells transiently co-transfected with the indicated GTPase pairs and treated with **1** (2 μM, 4 h). Each row shows NBD signal (green), the two GTPases, and merge, demonstrating co-occupancy of the same whorl structures. (Scale bar = 10 μm) (**F**) Representative CLSM images of HeLa cells transiently co-transfected with ARF6-EGFP (green) and Cytb5 (magenta) and treated with **1** (2 μM, 4 h), showing that Cytb5-marked ER whorls fail to recruit ARF6, in contrast to the ARF6-positive Golgi whorls induced by **1**. (Scale bar = 10 μm) (**G**) Representative CLSM images of HeLa cells treated with **1** (1 μM, 4 h) and loaded with fluorescent dextran, showing that dextran (magenta) is excluded from NBD-positive whorls (green), arguing against an endosomal origin for these structures. (Scale bar = 10 μm) (**H**) Representative CLSM images (merge) of A431, H460, HepG2, PANC-1, SH-SY5Y, and T98G cells transiently transfected with mCh-Rab11a and treated with **1** (2 μM, 4 h), showing that Rab11a recruitment to whorls occurs across multiple cell types. (Scale bar = 10 μm)

Whorls did not recruit membrane-associated proteins indiscriminately. Giantin and GalT retained their Golgi distribution, while Sec61β and calnexin (CANX) remained in the ER proteins; all four showed little whorl association (Fig. 2C). KDEL-containing proteins, KDELR2, Sec23a, GALNT2, TGN46, clathrin light chain A, the GPI-anchored protein CD59, and the palmitoylated tetraspanin CD9 also remained largely excluded (Fig. 2D; Figs. S11, S12).

We quantified recruitment by measuring whorl-to-cytosolic fluorescence enrichment and the fraction of whorls positive for each protein (Fig. 2D). Every tested GTPase showed substantially greater enrichment and occupancy than the resident, coat-associated, GPI-anchored, and non-GTPase palmitoylated proteins. The fatty-acid-handling enzyme ACSL3 also accumulated on whorls (Fig. S11), indicating that the selectivity extends beyond GTPases to at least some proteins involved in local fatty-acid metabolism. Both constitutively active and dominant-negative Rab11a mutants associated with whorls (Fig. S13), showing that recruitment does not require a single Rab11a nucleotide state.

Pairwise coexpression showed that ARF6 or CDC42 and Rab11a occupied the same individual whorls (Fig. 2E); the structures therefore do not segregate GTPases by family. Additional GTPase pairs, and coexpression with GalT-BFP, showed the same pattern (Fig. S14). Whorl origin, however, strongly affected recruitment. ER whorls induced by cytochrome b5 overexpression (*23*) failed to recruit ARF6 (Fig. 2F), and thapsigargin-induced ER whorls (*37*) showed the same result (Fig. S15). Fluorescent dextran also remained outside the whorl membranes despite their enrichment in Rab-family GTPases (Fig. 2G), arguing against an endosomal origin. Thus, Golgi-derived whorls recruit small GTPases through properties beyond multilamellar membrane geometry or endosomal identity.

Together, these results identify Golgi whorls as membrane compartments that override normal organelle boundaries and co-sequester small GTPases while excluding most tested resident, coat-associated, GPI-anchored, and palmitoylated proteins. This pattern points to shared membrane-association features, rather than native organelle identity or participation in one trafficking pathway, as the basis for whorl recruitment.

### Intact lipid anchors direct GTPase to cholesterol-poor Golgi whorls

To test whether shared membrane-anchoring features explain the unexpected co-recruitment, we disrupted covalent lipid anchors. Replacing the N-terminal glycine of ARF6 with alanine prevented N-myristoylation and markedly reduced ARF6 accumulation on whorls (Fig. 3A and 3B). Mutating the two C-terminal cysteines required for Rab11a geranylgeranylation likewise reduced recruitment (Fig. 3C, D). An ARF1 G2A mutant accumulated poorly on whorls (Fig. S16), and an N-myristoyltransferase inhibitor reduced ARF6 recruitment (Fig. 3E and Fig. S17). By contrast, a fluorescent CAAX construct did not accumulate on whorls (Fig. S18). Thus, intact lipid anchors promote ARF and Rab recruitment, but prenylation alone is insufficient; adjacent basic or amphipathic sequences and compatible membrane likely complete the targeting code.

**Figure 3.**
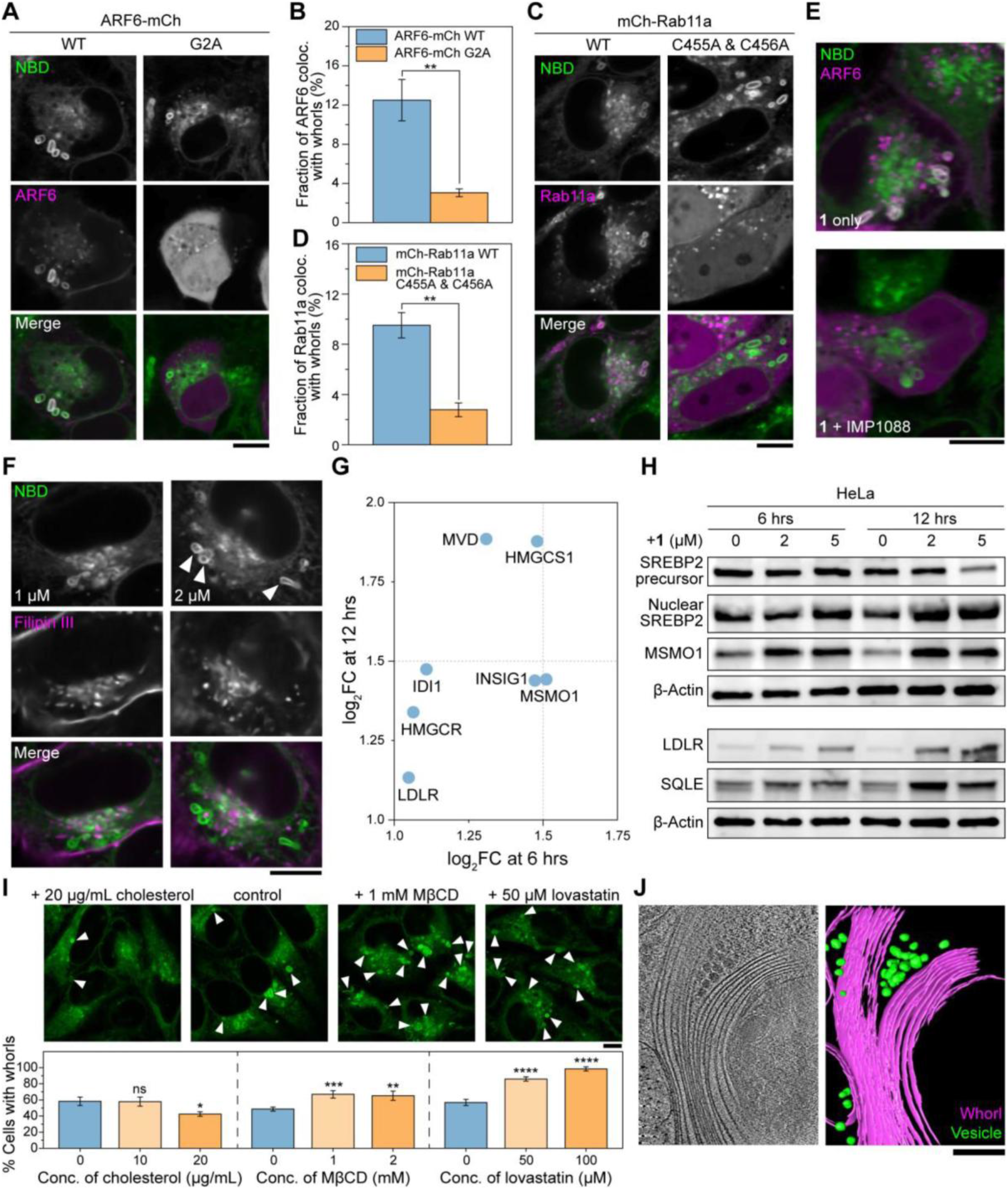
Intact lipid anchors promote GTPase partitioning into cholesterol-poor Golgi whorls. (**A**) Representative CLSM images of HeLa cells transiently transfected with ARF6-mCh (WT) or ARF6-mCh G2A and treated with **1** (2 μM, 4 h). (Scale bar = 10 μm) (**B**) Quantification of the fraction of ARF6 signal colocalized with whorls in HeLa cells transfected with ARF6-mCh WT or ARF6-mCh G2A and treated with **1** (2 μM, 4 h). (**C**) Representative CLSM images of HeLa cells transiently transfected with mCh-Rab11a (WT) or mCh-Rab11a C455A/C456A and treated with **1** (2 μM, 4 h). (Scale bar = 10 μm) (**D**) Quantification of the fraction of Rab11a signal colocalized with whorls in HeLa cells transfected with mCh-Rab11a WT or mCh-Rab11a C455A/C456A and treated with **1** (2 μM, 4 h). (**E**) Representative CLSM images of HeLa cells transiently transfected with ARF6-mCh and treated with **1** (2 μM, 4 h) alone (top) or co-treated with the N-myristoyltransferase inhibitor IMP-1088 (bottom). (Scale bar = 10 μm) (**F**) Representative CLSM images of HeLa cells treated with **1** at 1 μM (top) or 2 μM (bottom) for 4 h, fixed, and stained with filipin III. White arrowheads indicate whorls. (Scale bar = 10 μm) (**G**) Scatter plot of log2 fold changes in cholesterol biosynthesis and sterol homeostasis genes in HeLa cells treated with **1** at 6 h versus 12 h, relative to vehicle control. (**H**) Western blot analysis of SREBP-2 precursor, nuclear (cleaved) SREBP-2, and MSMO1 (top gel), and LDLR and SQLE (bottom gel), in HeLa cells treated with **1** at the indicated concentrations (0, 2, 5 μM) for 6 h or 12 h. β-Actin was used as a loading control for each gel. (**I**) Top: Representative CLSM images of HeLa cells treated with **1** (2 μM, 4 h) alone (control) or co-treated with cholesterol (20 μg/mL), MβCD (1 mM), or lovastatin (50 μM). For MβCD and lovastatin, cells were pretreated for 30 min before co-treatment with **1**; for cholesterol supplementation, cells were pretreated and co-treated with water-soluble cholesterol. White arrowheads indicate whorls. (Scale bar = 10 μm) Bottom: Quantification of the percentage of cells containing whorls under the conditions shown above. (**J**) Representative cryo-electron tomogram slice (left) and segmentation (right) of HeLa cells treated with **1** (10 μM, 4 h), showing densely packed concentric multilamellar membrane assemblies. Whorl membranes are shown in magenta and surrounding NBD-positive vesicles in green. (Scale bar = 200 nm) Data in (B, D, I) are presented as mean ± s.d. *P < 0.05, **P < 0.01, ***P < 0.001, ****P < 0.0001; ns, not significant (Student’s t-test). n = 3 independent cells per condition in B and D. n = 4 independent confocal views per condition in I.

We next asked whether whorls create a membrane environment that favors these lipid-anchored GTPases. We mapped accessible cholesterol with filipin III (*38*). Filipin strongly labeled the small NBD-positive puncta but weakly labeled mature whorl membranes (Fig. 3F). At 1 μM of **1**, filipin-positive puncta remained prominent; at 2 μM of **1**, filipin continued to label adjacent puncta but not the whorl rims. Compound **1** therefore segregated Golgi-derived membranes into filipin-poor whorls and filipin-retaining puncta. Because filipin detects accessible unesterified cholesterol rather than total membrane cholesterol, we operationally refer to the whorls as cholesterol poor.

Compound **1** also activated a sterol-homeostatic response. RNA-seq analysis at 6 h and 12 h showed coordinated induction of genes involved in cholesterol synthesis and uptake, including HMGCR, HMGCS1, MVD, IDI1, MSMO1, INSIG1, and LDLR (Fig. 3G and Fig. S19). Immunoblotting confirmed accumulation of nuclear SREBP2 and increased expression of MSMO1, SQLE, and LDLR (Fig. 3H). These responses show that cells sense reduced cholesterol availability or disrupted intracellular cholesterol distribution and activate the SREBP2 program.

Directly changing cholesterol availability shifted whorl formation in the predicted directions. Lovastatin inhibited de novo cholesterol synthesis and increased the percentage of cells containing whorls. Acute cholesterol extraction with methyl-β-cyclodextrin (MβCD) also increased whorl formation, whereas exogenous cholesterol supplementation reduced it (Fig. 3I). These complementary perturbations show that reduced cholesterol availability promotes whorl biogenesis.

Cryo-electron tomography resolved the structures as densely packed concentric multilamellar membrane lamellae (Fig. 3J), confirming the multilamellar architecture inferred from confocal imaging. Fluorescence recovery after photobleaching (FRAP) recovered incompletely, leaving an immobile NBD fraction of approximately 42% (Fig. S20); compound **1** therefore moved only partially within the whorl membranes. Together, these results define Golgi whorls as cholesterol-poor, multilamellar membrane assemblies that partition lipid-modified small GTPases.

### Distinct perturbations converge on GTPase-positive Golgi whorls

To test whether Golgi whorls represent a general response to lipid imbalance rather than a compound **1**-specific product, we examined three chemically distinct perturbations. CIL56 and tegavivint induce palmitoyl-CoA-dependent, nonapoptotic cell death and have been associated with Golgi dilation (*14, 29*), whereas the oxysterol 7-ketocholesterol (7-KC) perturbs cholesterol metabolism and can induce multilamellar membrane structures and nonapoptotic toxicity (*30, 31*). In HeLa cells expressing ARF6-mCherry, CIL56 and tegavivint generated ARF6-positive whorls within 4 h, whereas 7-KC produced similar structures after 16 h (Fig. 4A). Thus, distinct lipid perturbations converge on the same unexpected morphologically with different kinetics.

**Figure 4.**
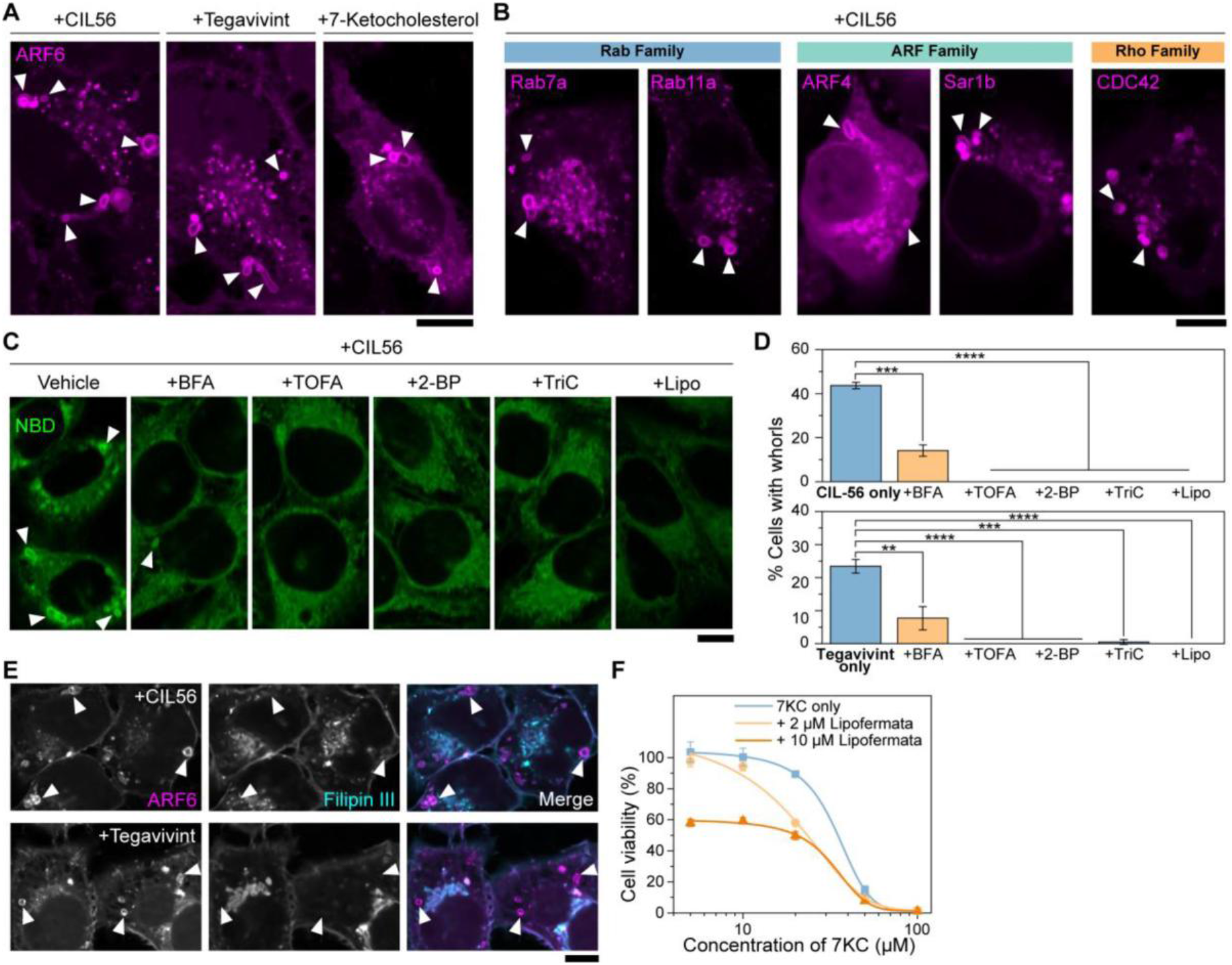
Distinct lipid perturbations converge on GTPase-positive Golgi whorls. (A) Representative CLSM images of a HeLa cell line stably expressing ARF6-mCh, treated with CIL56 (1 μM, 4 h), tegavivint (5 μM, 4 h), or 7-ketocholesterol (7-KC, 50 μM, 16 h). White arrowheads indicate whorls. (Scale bar = 10 μm) (B) Representative CLSM images of HeLa cells transiently transfected with RFP-tagged Rab7a, Rab11a (Rab family), ARF4, Sar1b (ARF family), or CDC42 (Rho family) and treated with CIL56 (1 μM, 4 h). White arrowheads indicate whorls. (Scale bar = 10 μm) (C) Representative CLSM images of HeLa cells stained with NBD-C6-ceramide and treated with CIL56 (1 μM, 4 h) alone (Vehicle) or co-treated with Brefeldin A (BFA, 50 nM), the acetyl-CoA carboxylase inhibitor TOFA (0.5 μM), the S-acylation inhibitor 2-BP (10 μM), the long-chain fatty acid activation inhibitor triacsin C (TriC, 0.5 μM), or the fatty acid uptake inhibitor lipofermata (Lipo, 10 μM). Cells were pretreated with each inhibitor for 30 min before co-treatment with CIL56 for 4 h. White arrowheads indicate whorls. (Scale bar = 10 μm) (D) Quantification of the percentage of cells containing whorls in HeLa cells treated with CIL56 (1 μM, top) or tegavivint (5 μM, bottom) alone or co-treated with the indicated inhibitors as in (C). (E) Representative CLSM images of the ARF6-mCh HeLa line treated with CIL56 (1 μM, 4 h, top) or tegavivint (5 μM, 4 h, bottom), fixed, and stained with filipin III. White arrowheads indicate whorls. (Scale bar = 10 μm) (F) Cell viability of HeLa cells treated with 7-KC at the indicated concentrations for 24 h, alone or co-treated with lipofermata (2 or 10 μM). Data in (D, F) are presented as mean ± s.d. **P < 0.01, ***P < 0.001, ****P < 0.0001 (Student’s t-test). n = 3 independent cells/experiments per condition.

CIL56-induced whorls recruited Rab7a, Rab11a, ARF4, Sar1b, and CDC42 (Fig. 4B), reproducing the cross-family GTPase recruitment observed with compound **1**. Tegavivint-induced whorls recruited a similar range of GTPases (Fig. S21). BFA, 2-BP, triacsin C, TOFA, and lipofermata each reduced whorl formation induced by CIL56 or tegavivint (Fig. 4C, D and Fig. S22). This shared inhibitor profile implicates Golgi integrity, fatty-acid metabolism, and S-acylation in all three responses. Because the compounds have pleiotropic actions, these data establish convergence on related lipid-dependent processes rather than an identical molecular pathway.

Filipin labeled CIL56- and tegavivint-induced ARF6-positive whorls weakly and remained concentrated in adjacent structures (Fig. 4E), matching the spatial segregation observed with compound **1**. CIL56 and tegavivint therefore also generate filipin-poor Golgi whorls.

7-KC reached a similar morphology through a different metabolic response. Inhibitors of S-acylation, acetyl-CoA carboxylase, or long-chain acyl-CoA synthesis did not protect cells from 7-KC-induced loss of viability (Fig. S23). Lipofermata instead enhanced 7-KC toxicity (Fig. 4F), consistent with a protective role for fatty-acid uptake and oxysterol esterification (*39*). These results separate the 7-KC response metabolically from the responses to compound **1**, CIL56, and tegavivint, although the mechanism that generates ARF6-positive whorls remains unresolved.

Together, these experiments show that several structurally distinct lipid perturbations converge on morphologically similar, GTPase-positive Golgi whorls. Compound **1**, CIL56, and tegavivint share dependencies on fatty-acid metabolism and S-acylation, whereas 7-KC reaches the same morphology through a distinct route. Golgi whorl formation therefore represents a recurring response to severe lipid imbalance rather than a compound **1**-specific effect.

### Golgi whorls disrupt membrane trafficking and trigger a poorly reversible, noncanonical loss of viability

Because Golgi whorls sequestered multiple trafficking GTPases, we tested whether cells retained normal membrane transport. Treatment with **1** moved the transferrin receptor (TfR) from the plasma membrane to perinuclear puncta in a concentration-dependent manner (Fig. 5A, B and S24). In a fluorescent transferrin pulse-chase assay, treated cells retained internalized transferrin and recycled it more slowly than vehicle-treated cells (Fig. 5C, Fig.S25). Compound **1** also moved EGFR from the plasma membrane to intracellular puncta in A431 cells (Fig. S26), showing that the trafficking defect extended beyond TfR.

**Figure 5.**
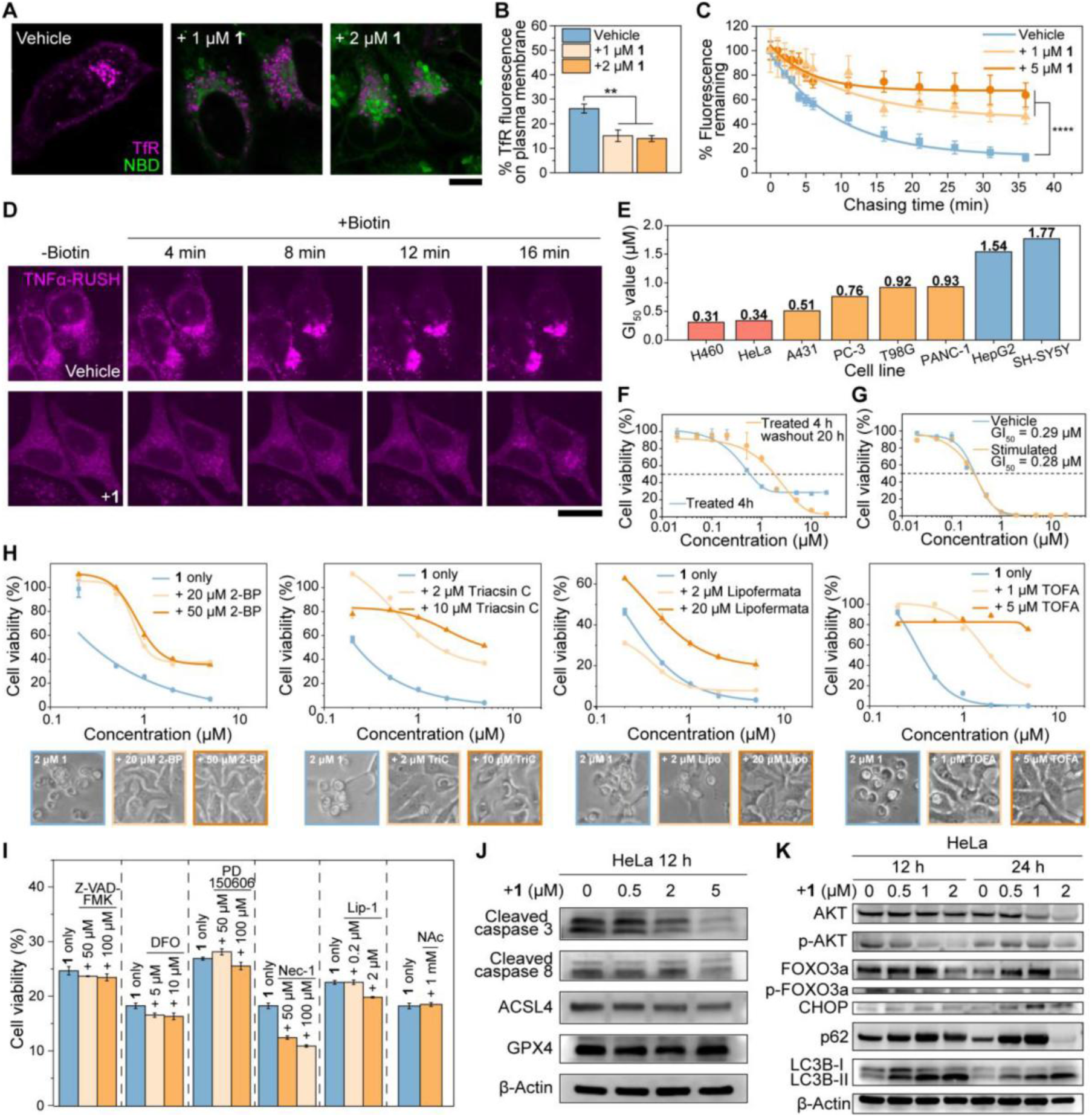
Golgi whorls disrupt membrane trafficking and trigger a poorly reversible, noncanonical loss of viability. (**A**) Representative CLSM images of HeLa cells transiently transfected with mCherry-TfR (magenta) and treated with **1** (0, 1, or 2 μM, 4 h), with NBD signal in green. Vehicle-treated cells show plasma membrane TfR, whereas **1**-treated cells show TfR redistribution to perinuclear regions. (Scale bar = 10 μm) (**B**) Quantification of the percentage of TfR fluorescence at the plasma membrane in HeLa cells treated with **1** at the indicated concentrations. (**C**) Fluorescent transferrin pulse-chase recycling assay in HeLa cells treated with **1** at the indicated concentrations. Percentage of internalized transferrin fluorescence remaining is plotted against chase time. (**D**) Representative time-lapse CLSM images of a RUSH assay in HeLa cells expressing TNFα-mCherry, treated with vehicle (top) or **1** (2 μM, bottom). Anterograde transport was initiated with biotin, and images were acquired at the indicated times. (Scale bar = 20 μm) (**E**) GI_50_ values of **1** across a panel of cell lines determined by cell viability assay. (**F**) Cell viability of HeLa cells treated with **1** for 4 h then either maintained or washed out for 20 h, plotted against concentration of **1**, showing a point of no return above which washout fails to rescue viability. (**G**) Cell viability of parental HeLa cells or cells subjected to prolonged exposure to **1** over 4 weeks (“stimulated”), plotted against concentration of **1**. The comparable GI_50_ values (0.29 μM versus 0.28 μM) indicate that long-term exposure does not induce acquired resistance. (**H**) Cell viability of HeLa cells treated with **1** alone or co-treated with the indicated lipid-pathway inhibitors, plotted against concentration of **1**. Inhibitors target S-acylation (2-BP), long-chain fatty acid activation (triacsin C), fatty acid uptake (lipofermata), and acetyl-CoA carboxylase (TOFA). Representative bright-field images of cells treated with **1** (2 μM) alone or with each inhibitor are shown below. (**I**) Cell viability of HeLa cells treated with **1** alone or co-treated with inhibitors of apoptosis (Z-VAD-FMK), ferroptosis (DFO, Lip-1), calpain activity (PD 150606), necroptosis (Nec-1), or oxidative stress (NAc), at the indicated concentrations. (**J**) Western blot analysis of cleaved caspase-3, cleaved caspase-8, ACSL4, and GPX4 in HeLa cells treated with **1** at the indicated concentrations (0, 0.5, 2, 5 μM) for 12 h. β-Actin was used as a loading control. (**K**) Western blot analysis of AKT, p-AKT, FOXO3a, p-FOXO3a, CHOP, p62, and LC3B-I/LC3B-II in HeLa cells treated with **1** at the indicated concentrations (0, 0.5, 1, 2 μM) for 12 h or 24 h. β-Actin was used as a loading control. Data in (B, C, I) are presented as mean ± s.d. GI_50_ values in (E, F, G, H) were determined by nonlinear regression. **P < 0.01, ****P < 0.0001 (Student’s t-test). n = 5 independent cells per condition in B and C. n = 3 independent experiments per condition in (I).

Compound **1** also slowed anterograde transport. In RUSH assays, whorl-containing cells moved TNFα-mCherry and E-cadherin-mCherry from the ER towards the plasma membrane more slowly than control cells after biotin release (Fig. 5D and Fig. S27). Phalloidin staining revealed more F-actin and thickened stress fibers in whorl-containing cells (Fig. S28), consistent with disrupted Rho-family GTPases signaling. Untargeted lipidomic detected increased lysolipids and transient changes in major phospholipid classes (Fig. S29). Thus, whorl accumulation disrupted endocytic recycling and anterograde transport while remodeling the cytoskeleton and lipid composition.

Compound **1** reduced viability across several cell lines, and H460 cells showed the lowest GI_50_ in the panel (Fig. 5E). A 4-h exposure followed by a 20-h washout restored viability at lower concentrations but not at concentrations of 2 μM or higher (Fig. 5F). This threshold coincided with near-maximal whorl prevalence (Fig. 1E), linking extensive whorl accumulation to a poorly reversible loss of viability over the experimental interval. HeLa cells maintained in the presence of **1** for four weeks remained as sensitive as parental cells and retained similar GI_50_ values (Fig. 5G), so prolonged exposure did not produce detectable resistance under these culture conditions. Total ARF6 abundance remained unchanged (Fig. S30), excluding altered ARF6 expression as an explanation for the maintained sensitivity.

Lipid-metabolism inhibitors that suppressed whorl formation also improved cell viability during treatment with compound **1** (Fig. 5H and S31). Peptide analogs lacking the thioester warhead (**2** and **3**) formed no whorls and caused less cytotoxicity (Figs. S32-S34). These matching structure–activity and rescue relationships link the loss of viability to S-palmitoylation-dependent whorl formation rather than to nonspecific exposure to the peptide scaffold.

Inhibitors of caspases, ferroptosis, calpain activity, necroptosis, or oxidative stress failed to substantially rescue viability under the tested conditions (Fig. 5I). ULK1 inhibition instead enhanced the effect of compound **1** (Fig. S35), suggesting that ULK1-dependent stress responses provide partial protection. These pharmacological results argue against a dominant role for the tested canonical death pathways, although they do not exclude secondary contributions.

Immunoblotting detected no increase in cleaved caspase-3 or caspase-8 and no substantial change in GPX4 or ACSL4 abundance (Fig. 5J), providing no biochemical evidence for robust apoptosis or ferroptosis under these conditions. Compound **1** reduced AKT and FOXO3a phosphorylation and increased CHOP, p62, and LC3B-I abundance (Fig. 5K). These changes link whorl accumulation to reduced survival signaling, ER stress, and altered autophagy-related protein turnover. Direct flux assays will be needed to determine whether cells also block autophagic degradation.

Together, these results show that Golgi whorls disrupt multiple membrane-trafficking routes and accompany a poorly reversible loss of viability. Reduced AKT–FOXO3a signaling, ER stress, and altered autophagy markers characterize the response, but inhibitors of several canonical death pathways do not prevent it. We therefore describe the phenotype as noncanonical while leaving its formal cell-death classification unresolved.

## Discussion

Two prevailing expectations framed this study. First, compartment-specific regulators and lipid cues should keep ARF-, Rab-, and Rho-family GTPases segregated among their native membranes. Indeed, super-resolution imaging of endogenously tagged ARF GTPases showed that even closely related ARF paralogs occupy distinct nanoscale domains across the Golgi, ERGIC, and TGN (*40*). Second, stress-induced multilamellar whorls should behave as passive or nonspecific membrane accumulations. Our results violate both expectations. The Golgi-derived whorls enriched diverse small GTPases while excluding most tested transmembrane residents, coat-associated proteins, a GPI-anchored protein, and other palmitoylated proteins. Previous studies identified them primarily in the ER or lysosomal system during protein overload, lipid-storage disorders, drug toxicity, or altered autophagy (*23-26*). Yet cytochrome b5-induced ER whorls did not recruit ARF6 (Fig. 2F), nor did thapsigargin-induced ER whorls (Fig. S15), and the Golgi whorls lacked LAMP1 and ATG14 (Fig. S11, S12). Multilamellar geometry and generalized stress therefore cannot explain the selective partitioning. The induction of related GTPase-positive whorls by CIL56, tegavivint, and 7-KC further argues that the phenotype reflects recurring response to severe lipid imbalance rather than peptide aggregation.

The evidence instead supports a membrane-state model that reverses the usual causal picture of organelle identity. Small GTPases normally help organize membranes, but here an altered membrane state reorganizes the GTPases. We use “jamsome,” introduced above, as an operational term for these induced Golgi-derived, multilamellar membrane sinks because they co-sequester trafficking regulators and jam membrane traffic. We use the term descriptively, not to designate a constitutive organelle class. In the proposed model (Fig. 6), excessive palmitoylated peptide accumulates in discrete Golgi regions, reduced cholesterol availability promotes compositional segregation, and the filipin-poor membrane curls into a multilamellar whorl. Compatible lipid anchors and adjacent targeting sequences then partition multiple GTPase families into this membrane state, whereas cholesterol-retaining puncta preserve many Golgi residents. The failure of a generic CAAX construct to enter the whorls shows that a lipid anchor is necessary but not sufficient. ARF-, Rab-, and Sar1-family proteins can also deform, constrict, or tether membranes (*41-43*), raising the possibility that recruited GTPases reinforce jamsome growth in a feed-forward loop. The present experiments do not directly test that step.

**Figure 6.**
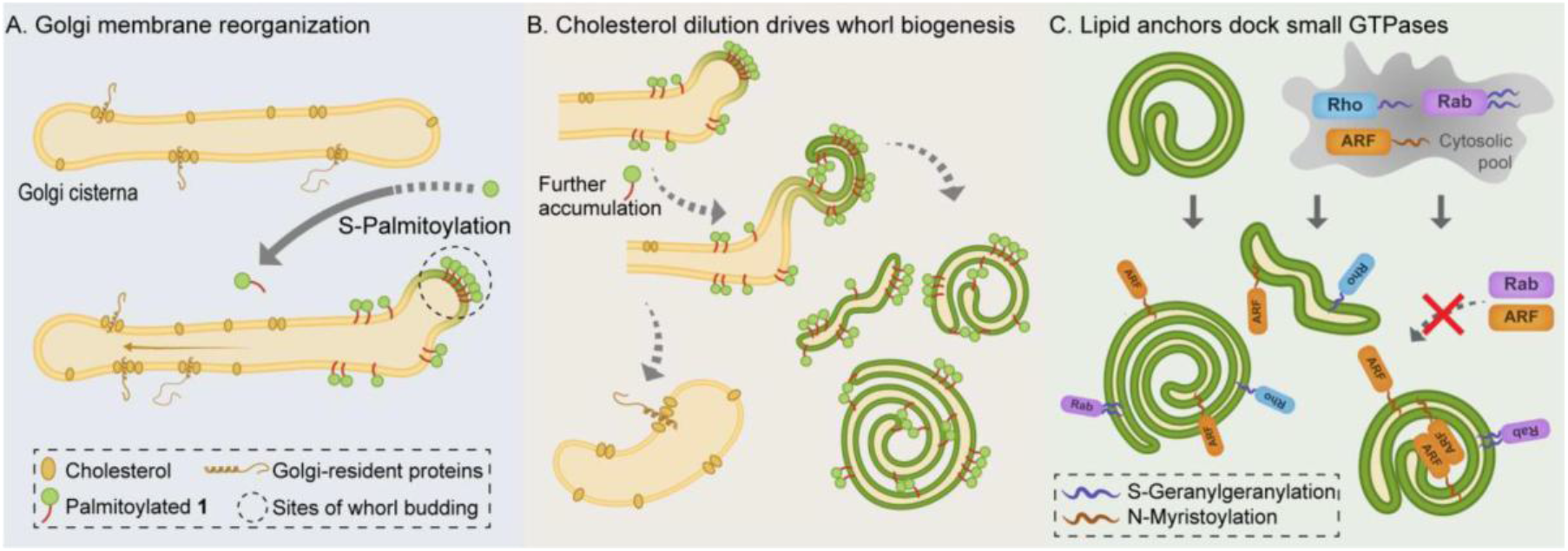
Proposed model. Severe Golgi lipid imbalance creates a selective multilamellar membrane sink that we term a “jamsome”. (A) Palmitoylated **1** (green) accumulates in discrete Golgi membrane regions and promotes compositional segregation. These regions lose filipin-detectable cholesterol, whereas adjacent Golgi-derived puncta retain cholesterol and resident proteins. (B) Continued accumulation of palmitoylated **1** curls the cholesterol-poor membrane into a multilamellar whorl. (C) Compatible combinations of lipid anchors, amphipathic elements, and adjacent targeting sequences partition ARF-, Rab-, and Rho-family GTPases into the whorl, whereas most tested resident and coat-associated proteins remain excluded. The resulting jamsome co-sequesters trafficking regulators, slows endocytic recycling and anterograde transport, and promotes cellular stress. The dashed causal link from GTPase accumulation to further membrane curling represents a testable feed-forward hypothesis.

Independent observations converge on this interpretation. Compound **1** first accumulates with GalT and Giantin, live imaging captures whorls emerging from Golgi-associated membranes, BFA suppresses their formation, and experimentally induced ER whorls behave differently. The Golgi may be especially susceptible because it concentrates ZDHHC acyltransferases and continuously sorts sterols, fatty acids, and membrane cargo (*17-20*). Bidirectional cholesterol manipulations further connect reduced sterol availability to whorl biogenesis: lowering cholesterol promotes the phenotype, whereas supplementation suppresses it (Fig. 3I and Fig. S36). Lipid-anchor mutations reduce GTPase capture, cryo-electron tomography confirms the multilamellar architecture, and trafficking assays link jamsome accumulation to impaired recycling and anterograde transport. These distinct perturbations, imaging modalities, and functional readouts provide consilience for the central update: severe lipid imbalance can create an organized membrane sink that overrides native organelle targeting. Other organelles nevertheless participate in the cellular response. Sar1b recruitment (Fig. 2A), the effect of salubrinal (Fig. S37), and peripheral whorls at later times (Fig. S38) suggest communication with ER stress and possibly secondary membrane remodeling. Direct organelle-lineage tracing will be needed to establish whether all late structures remain Golgi-derived.

Several limitations define the current mechanistic interpretation. Filipin reports accessible unesterified cholesterol rather than total cholesterol or membrane order, so the data establish a filipin-poor state but not a formally defined membrane phase. The anchor mutations demonstrate a requirement for myristoylation and geranylgeranylation in representative proteins but do not establish direct anchor-lipid binding. The shared inhibitor profiles of compound **1**, CIL56, and tegavivint indicate convergence on lipid-dependent processes without proving an identical upstream pathway, and 7-KC may reach a similar morphology through a distinct route. Finally, the current data separate the viability phenotype from several canonical death programs but do not identify its execution mechanism. The jamsome model makes falsifiable predictions: interventions that block GTPase capture without preventing whorl formation should uncouple membrane morphology from trafficking failure; conversely, strengthening a compatible composite membrane-targeting code should enhance sequestration. Direct sterol measurements, membrane reconstitution, organelle-lineage tracing, and selective depletion of recruited GTPases can test these predictions.

## Conclusion

The central result is not simply that lipid imbalance damages the Golgi. It is that lipid imbalance creates a selective membrane state that overrides normal organelle boundaries. We propose calling these induced Golgi-derived membrane sinks jamsomes. Jamsomes segregate from cholesterol-retaining Golgi fragments, co-sequester ARF-, Rab-, and Rho-family GTPases through composite membrane-targeting features, and progressively interrupt endocytic recycling and anterograde transport. Multiple chemically distinct perturbations generate related structures, indicating that jamsome formation is a recurring membrane-remodeling response rather than a compound-specific artifact.

These findings change the causal relationship between membrane identity and trafficking regulators. Small GTPases do not only organize organelles; under severe lipid imbalance, membrane composition and architecture can reorganize small GTPases across their native compartments. The jamsome model therefore connects Golgi cholesterol availability, multilamellar remodeling, GTPase partitioning, trafficking failure, stress signaling, and cell fate in one testable framework. Future work should determine whether jamsomes arise in physiological or disease contexts, whether GTPase clustering actively drives their growth, and whether preventing selective sequestration preserves trafficking and viability. Establishing those points would determine whether jamsomes represent an experimentally induced state or a broader cellular response to lipid imbalance.

## Acknowledgments

We thank T. Kirchhausen, L. Lu, J. Liou and J. Baskin for their insightful discussions and suggestions. TEM samples were prepared and imaged at the Brandeis Electron Microscopy Facility. We thank the Brandeis Light Microscopy Facility (RRID:SCR_025892) for assistance. Untargeted lipidomics profiling was conducted at the BIDMC-Harvard Mass Spectrometry Facility. RNA-seq analysis was conducted at Azenta/GENEWIZ. The authors acknowledge the use of ChatGPT to assist with drafting and language refinement.

## Funding

National Institutes of Health grant CA142746 (BX)

National Science Foundation DMR-2011846 (BX)

## Author contributions

Conceptualization: QZ, WT, BX

Methodology: QZ, WT, BX

Investigation: QZ, IAR, JH, SJB, JZ, BX

Visualization: QZ, BX

Funding acquisition: BX

Project administration: BX

Supervision: BX

Writing – original draft: QZ, BX

Writing – review & editing: QZ, JLS, BX

## Competing interests

Authors declare that they have no competing interests.

## Data and materials availability

All data are available in the main text or the supplementary materials.

## Supplementary Materials

Materials and Methods

Supplementary Text

Figs. S1 to S38

## Notes

### Competing Interest Statement

The authors have declared no competing interest.

## References and Notes

1. W. D. Heo et al., PI(3,4,5)P3 and PI(4,5)P2 lipids target proteins with polybasic clusters to the plasma membrane. Science 314, 1458–1461 (2006).

2. S. M. F et al., Mechanisms and functions of protein S-acylation. Nat Rev Mol Cell Biol 25, 488–509 (2024).

3. P. Chlanda et al., Palmitoylation Contributes to Membrane Curvature in Influenza A Virus Assembly and Hemagglutinin-Mediated Membrane Fusion. J Virol 91, (2017).

4. A. M. Ernst et al., S-Palmitoylation Sorts Membrane Cargo for Anterograde Transport in the Golgi. Dev Cell 47, 479–493.e477 (2018).

5. P. Song et al., Palmitic acid and palmitoylation in cancer: Understanding, insights, and challenges. Innovation (Camb) 6, 100918 (2025).

6. E. W. Tate, L. Soday, A. L. de la Lastra, M. Wang, H. Lin, Protein lipidation in cancer: mechanisms, dysregulation and emerging drug targets. Nature Reviews Cancer 24, 240–260 (2024).

7. M. Yeste-Velasco, M. E. Linder, Y. J. Lu, Protein S-palmitoylation and cancer. Biochim. Biophys. Acta Rev. Cancer 1856, 107–120 (2015).

8. A. Yanai et al., Palmitoylation of huntingtin by HIP14 is essential for its trafficking and function. Nat Neurosci 9, 824–831 (2006).

9. M. Fukata, Y. Fukata, H. Adesnik, R. A. Nicoll, D. S. Bredt, Identification of PSD-95 palmitoylating enzymes. Neuron 44, 987–996 (2004).

10. K. Tabata et al., Palmitoylation of ULK1 by ZDHHC13 plays a crucial role in autophagy. Nature Communications 15, 7194 (2024).

11. P. Fontana et al., Small-molecule GSDMD agonism in tumors stimulates antitumor immunity without toxicity. Cell 187, 6165–6181.e6122 (2024).

12. A. Balasubramanian et al., The palmitoylation of gasdermin D directs its membrane translocation and pore formation during pyroptosis. Science Immunology 9, eadn1452.

13. B. Huang et al., Palmitoylation-dependent regulation of GPX4 suppresses ferroptosis. Nat Commun 16, 867 (2025).

14. P. J. Ko et al., A ZDHHC5-GOLGA7 Protein Acyltransferase Complex Promotes Nonapoptotic Cell Death. Cell Chem Biol 26, 1716–1724.e1719 (2019).

15. N. Zhang et al., Palmitoylation licenses RIPK1 kinase activity and cytotoxicity in the TNF pathway. Mol Cell 84, 4419–4435 e4410 (2024).

16. W. Tan et al., Cycling molecular assemblies for Golgi imaging and disruption. Nat Commun, (2026).

17. O. Rocks et al., The palmitoylation machinery is a spatially organizing system for peripheral membrane proteins. Cell 141, 458–471 (2010).

18. J. T. Swarthout et al., DHHC9 and GCP16 constitute a human protein fatty acyltransferase with specificity for H- and N-Ras. Journal of Biological Chemistry 280, 31141–31148 (2005).

19. C. Salaün, C. Locatelli, F. Zmuda, J. C. González, L. H. Chamberlain, Accessory proteins of the zDHHC family of S-acylation enzymes. Journal of Cell Science 133, (2020).

20. M. A. Kahlson et al., Functional dissection of the zDHHC palmitoyltransferase 5–golgin A7 palmitoylation complex. Journal of Biological Chemistry 301, (2025).

21. J. S. Goodwin et al., Depalmitoylated Ras traffics to and from the Golgi complex via a nonvesicular pathway. J Cell Biol 170, 261–272 (2005).

22. K. F. Ferri, G. Kroemer, Organelle-specific initiation of cell death pathways. Nature Cell Biol. 3, E255–263 (2001).

23. E. L. Snapp et al., Formation of stacked ER cisternae by low affinity protein interactions. Journal of Cell Biology 163, 257–269 (2003).

24. J. Guo et al., Neuronal ceroid lipofuscinosis in a German Shorthaired Pointer associated with a previously reported CLN8 nonsense variant. Mol. Genet. Metab. Rep. 21, (2019).

25. R. Kerkelä et al., Cardiotoxicity of the cancer therapeutic agent imatinib mesylate. Nat. Med. 12, 908–916 (2006).

26. S. Schuck, C. M. Gallagher, P. Walter, ER-phagy mediates selective degradation of endoplasmic reticulum independently of the core autophagy machinery. Journal of Cell Science 127, 4078–4088 (2014).

27. D. M. Yang, A. S. Chiang, Formation of a whorl-like autophagosome by Golgi apparatus engulfing a ribosome-containing vacuole in corpora allata of the cockroach Diploptera punctata. CELL TISSUE RES. 287, 385–391 (1997).

28. E. Bejarano, M. Cabrera, L. Vega, J. Hidalgo, A. Velasco, Golgi structural stability and biogenesis depend on associated PKA activity. Journal of Cell Science 119, 3764–3775 (2006).

29. L. Leak et al., Tegavivint triggers TECR-dependent nonapoptotic cancer cell death. Nature Chemical Biology, (2025).

30. K. Casazza, S. M. Cologna, E. Berry-Kravis, J. Jarnes-Utz, F. D. Porter, Biomarker Validation in NPC1: Foundations for Clinical Trials and Regulatory Alignment. J. Inherit. Metab. Dis. 48, (2025).

31. A. Véjux et al., Cytotoxic oxysterols induce caspase-independent myelin figure formation and caspase-dependent polar lipid accumulation. Histochem. Cell Biol. 127, 609–624 (2007).

32. W. Tan, Q. Zhang, P. Hong, B. Xu, Structure-Activity Relationship of Cycling Molecular Assemblies for Golgi Targeting and Disruption. J Med Chem, (2026).

33. P. Chardin, F. McCormick, Brefeldin A: The Advantage of Being Uncompetitive. Cell 97, 153–155 (1999).

34. J. B. McCabe, L. G. Berthiaume, N-terminal protein acylation confers localization to cholesterol, sphingolipid-enriched membranes but not to lipid rafts/caveolae. Mol Biol Cell 12, 3601–3617 (2001).

35. G. Kulakowski et al., Lipid packing defects and membrane charge control RAB GTPase recruitment. Traffic 19, 536–545 (2018).

36. M. G. t. Hanna et al., Sar1 GTPase Activity Is Regulated by Membrane Curvature. J Biol Chem 291, 1014–1027 (2016).

37. F. Xu et al., COPII mitigates ER stress by promoting formation of ER whorls. Cell Research 31, 141–156 (2021).

38. C. P. Muller et al., Filipin as a flow microfluorometry probe for cellular cholesterol. Cytometry 5, 42–54 (1984).

39. J. W. Lee, J.-D. Huang, I. R. Rodriguez, Extra-hepatic metabolism of 7-ketocholesterol occurs by esterification to fatty acids via cPLA2α and SOAT1 followed by selective efflux to HDL. Biochimica et Biophysica Acta (BBA) - Molecular and Cell Biology of Lipids 1851, 605–619 (2015).

40. L. Wong-Dilworth et al., STED imaging of endogenously tagged ARF GTPases reveals their distinct nanoscale localizations. Journal of Cell Biology 222, (2023).

41. X. Pang et al., Structural elucidation of how ARF small GTPases induce membrane tubulation for vesicle fission. Proc Natl Acad Sci U S A 122, e2417820122 (2025).

42. J. Mima, Self-assemblies of Rab- and Arf-family small GTPases on lipid bilayers in membrane tethering. Biophys. Rev. 13, 531–539 (2021).

43. K. R. Long et al., Sar1 assembly regulates membrane constriction and ER export. J Cell Biol 190, 115–128 (2010).

