## Supplemental information for "Lipid Imbalance Generates Golgi Whorls that Sequester Small GTPases"

**The PDF file includes:**

Materials and Methods
Supplementary Text
Figs. S1 to S38

**Materials and Methods**

Materials

2-Cl trityl chloride resin (1.0-1.2 mmol/g) and HBTU were purchased from GL Biochem (Shanghai, China). N, N-diisopropylethylamine (DIEA) and solvents were purchased from Fisher Scientific (Waltham, MA). β-alanine was purchased from Indofine Chemical Company (Hillsborough, NJ). Acetyl anhydride was purchased from Sigma-Aldrich (St. Louis, MO). Fmoc-D-4,4'-biphenylalanine was purchased from ChemScene (Monmouth Junction, NJ). 3-(*Tert*-butoxy)propanoic acid was purchased from A2B Chem (San Diego, CA). 4-Chloro-7-nitrobenzofurazan was purchased from Alfa Aesar (Ward Hill, MA). Acetyl chloride was purchased from TCI America (Portland, OR). 3-(Acetylthio)propionic acid was purchased from 1PlusChem (San Diego, CA). N-Boc-ethylenediamine was purchased from Ambeed (Buffalo Grove, IL). Brefeldin A (BFA), lipofermata, ML211, DC661, Necrostatin-1 (Nec-1), Liproxstatin-1 (Lip-1), SBI-0206965, bragsin1, NAV-2729, CIL56, tegavivint, lovastatin, IMP1088 salubrinal and Golgicide A were purchased from MedChemExpress (Monmouth Junction, NJ). 2-Bromohexadecanoic acid (2-BP) and transferrin Alexa Fluor™ 647 conjugate were purchased from Fisher Scientific (Waltham, MA). Triacsin C (TriC) and Filipin III were purchased from Cayman Chemical (Ann Arbor, MI). 5-(tetradecyloxy)-2-furancarboxylic acid (TOFA) and N-acetylcysteine (NAc) were purchased from Selleck Chemicals (Houston, TX). Z-VAD-FMK was purchased from APExBIO (Houston, TX). Deferoxamine mesylate (DFO) and 5-(N-ethyl-N-isopropyl)-amiloride (EIPA) were purchased from Santa Cruz Biotechnology (Dallas, TX). PD150606 was purchased from Tocris Bioscience (Minneapolis, MN). All the chemical reagents and solvents were used as received from commercial sources without further purification.

CellLight™ Golgi-RFP, BacMam 2.0 (Catalog #C10593) was purchased from Invitrogen (Waltham, MA). ARF6 monoclonal antibody (Catalog #5740), cleaved caspase-3 monoclonal antibody (Catalog #9664), cleaved caspase-8 monoclonal antibody (Catalog #98134), ACSL4 monoclonal antibody (Catalog #38493), GPX4 antibody (Catalog #52455), FOXO3a monoclonal antibody (Catalog #2497), phospho-FoxO3a (Ser318/321) antibody (Catalog #9465), BiP monoclonal antibody (Catalog #3177), SQSTM1/p62 antibody (Catalog #5114), PERK monoclonal antibody (Catalog #3192) and phospho-PERK (Thr980) monoclonal antibody (Catalog #3179) were purchased from Cell Signaling Technology (Danvers, MA). Raptor polyclonal antibody (Catalog #42-4000), mTor polyclonal antibody (Catalog #PA5-34663), goat antirabbit IgG (H+L) secondary antibody, HRP (Catalog #31460), goat anti-rabbit IgG (Heavy chain), superclonal™ recombinant secondary antibody, Alexa Fluor™ 647 (Catalog #A27040), SuperSignal™west pico PLUS chemiluminescent substrate (Catalog #34579), SuperBlock™ (TBS) blocking buffer (Catalog #37535), Restore™ PLUS western blot stripping buffer (Catalog #46428), and Halt™ protease inhibitor cocktail (100X) (Catalog #78430) were purchased from Invitrogen (Waltham, MA). AKT1 + AKT2 + AKT3 monoclonal antibody (Catalog #ab179463), AKT1 (pS473) + AKT2 (pS474) + AKT3 (pS472) monoclonal antibody (Catalog #ab192623), LC3B monoclonal antibody (Catalog #ab192890), beta Actin antibody (Catalog # ab8227) and goat anti-rabbit IgG H&L (Alexa Fluor® 647) (Catalog #ab150079) were purchased from abcam (Waltham, MA). CHOP polyclonal antibody (Catalog #15204-1-AP) was purchased from Proteintech (Rosemont, IL).

EGFR-mApple was a gift from the Thorsten Wohland lab (Addgene plasmid #203773). EndophilinA1-mRuby2 was a gift from the Cheng-Han Yu lab (Addgene plasmid #171945). GalToxBFP was a gift from the Erik Snapp lab (Addgene plasmid #68073). Lifeact-mCherry was a gift from the Klaus Hahn lab (Addgene plasmid #193300). mCh-Mfn2 was a gift from the Gia Voeltz lab (Addgene plasmid #141156). mCh-Rab5 was a gift from the Gia Voeltz lab (Addgene plasmid #49201). mCh-Rab7A was a gift from the Gia Voeltz lab (Addgene plasmid #61804). mCherry was a gift from the Rob Parton lab (Addgene plasmid #176016). mCherry-ATG14-C-18 was a gift from the Michael Davidson lab (Addgene plasmid #54989). mCherry-CaaX(Hras) was a gift from the Rob Parton lab (Addgene plasmid #108886). mCherry-Calnexin-N-14 was a gift from the Michael Davidson lab (Addgene plasmid #55005). mCherry-Caveolin-C-10 was a gift from the Michael Davidson lab (Addgene plasmid #55008). mCherry-CD9-10 was a gift from the Michael Davidson lab (Addgene plasmid #55013). mCherry-CDC42-C-10 was a gift from the Michael Davidson lab (Addgene plasmid #55014). mCherry-Cytb5 was a gift from the Uri Manor lab (Addgene plasmid #182579). mCherry-Rab11a-7 was a gift from the Michael Davidson lab (Addgene plasmid #55124). mCherry-Rab4a-7 was a gift from the Michael Davidson lab (Addgene plasmid #55125). mCherry-Sec23A was a gift from the Jennifer Lippincott-Schwartz lab (Addgene plasmid #166894). mCherry-Sec61b-C1 was a gift from the Jennifer Lippincott-Schwartz lab (Addgene plasmid #90994). mCherry-TFR-20 was a gift from the Michael Davidson lab (Addgene plasmid #55144). mCherry-TGNP-N-10 was a gift from the Michael Davidson lab (Addgene plasmid #55145). mCherry-TOMM20-N-10 was a gift from the Michael Davidson lab (Addgene plasmid #55146). mScarlet-Rab9a was a gift from the Gia Voeltz lab (Addgene plasmid #169074). pCAG/hArf4(WT)-mCherry was a gift from the Kazuhisa Nakayama lab (Addgene plasmid #79406). pCMV-dTomato-ORP9-PH was a gift from the Shinya Tsukiji lab (Addgene plasmid #214268). pcDNA3/hArf1(WT)-mCherry was a gift from the Kazuhisa Nakayama lab (Addgene plasmid #79419). pcDNA3/hArf3(WT)-mCherry was a gift from the Kazuhisa Nakayama lab (Addgene plasmid #79420). pcDNA3/hArf5(WT)-mCherry was a gift from the Kazuhisa Nakayama lab (Addgene plasmid #79421). pcDNA3/hArf6(WT)-mCherry was a gift from the Kazuhisa Nakayama lab (Addgene plasmid #79422). pCS2-CIB1-mCherry-Rab8A was a gift from the Heidi Hehnly lab (Addgene plasmid #194333). pCS2-mCherry-Rab11a was a gift from the Heidi Hehnly lab (Addgene plasmid #184035). pCS2-mCherry-Rab11a (Q70L) was a gift from the Heidi Hehnly lab (Addgene plasmid #184037). pCS2-mCherry-Rab11a (S25N) was a gift from the Heidi Hehnly lab (Addgene plasmid #184036). pEFIRES-P-ACSL3-mCherry was a gift from the Elina Ikonen lab (Addgene plasmid #87158). pEF.myc.ER-E2-Crimson was a gift from the Benjamin Glick lab (Addgene plasmid #38770). pEGFP-N3/hArf6(WT)-EGFP was a gift from the Kazuhisa Nakayama lab (Addgene plasmid #79423). pLAMP1-mCherry was a gift from the Amy Palmer lab (Addgene plasmid #45147). pME-mCherry-FLAG-CD59-GPI was a gift from the Reika Watanabe lab (Addgene plasmid #50378). pmCherry-LCa was a gift from the Tom Kirchhausen lab (Addgene plasmid #53972). pmCherry-N1-GalT was a gift from the Lei Lu lab (Addgene plasmid #87327). pmScarlet-HSP70 was a gift from the Vincent Timmerman lab (Addgene plasmid #163790). pmScarlet3-Giantin_C1 was a gift from the Dorus Gadella lab (Addgene plasmid #189773). pSIRV-AP-1-mCherry was a gift from the Peter Steinberger lab (Addgene plasmid #118095). pTag-BFP-C-h-Rab11a was a gift from the James Johnson lab (Addgene plasmid #79805). pTriEx-mCherry-PA-Rac1 was a gift from the Klaus Hahn lab (Addgene plasmid #22027). Sar1b-mOrange2 was a gift from the Jennifer Lippincott-Schwartz lab (Addgene plasmid #166899). Str-KDEL_SBP-mCherry-Ecadherin was a gift from the Franck Perez lab (Addgene plasmid #65287). Str-KDEL_TNF-SBP-mCherry was a gift from the Franck Perez lab (Addgene plasmid #65279). tdTomato-ELP-1-25 was a gift from the Michael Davidson lab (Addgene plasmid #58091). VAMP3-mCherry was a gift from the Geert van den Bogaart lab (Addgene plasmid #92423). VPS13A^mCherry was a gift from the Pietro De Camilli lab (Addgene plasmid #118758). Xfect™ Transfection Reagent (Catalog #631317) was purchased from TaKaRa.

Minimum Essential Medium (MEM), Dulbecco's Modified Eagle Medium (DMEM), Ham's F-12 Media (F-12), fetal bovine serum (FBS) and Penicillin-Streptomycin were purchased from Gibco by Life Technologies (Carlsbad, CA). RPMI-1640 medium and Eagle's Minimum Essential Medium (EMEM) were purchased from American Type Culture Collection (ATCC, Manassas, VA).

Instruments

All compounds were purified by a reverse phase HPLC (Agilent 1100 Series) equipped with an Xterra C18 RP column. HPLC grade acetonitrile (0.1% TFA) and HPLC grade water (0.1% TFA) were used as eluents. The LC-MS spectra were obtained with a Bruker Elute PLUS UHPLC with a Bruker timsTOF Pro. Transmission electron microscopic images were obtained on Morgagni 268 transmission electron microscope. Fluorescent images were taken by Nikon AX-R Confocal System at the lens of 60× with oil.

Methods

Synthesis of NBD-ethylenediamine

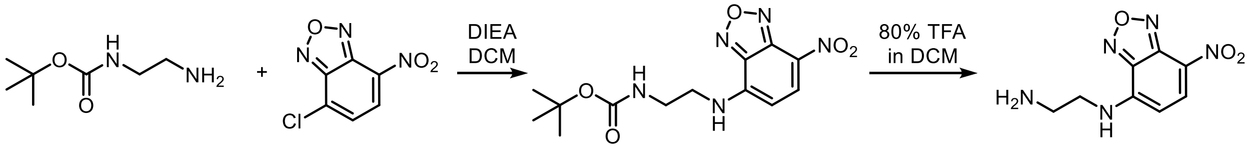


**Scheme S1.** Synthetic route of NBD-ethylenediamine.

N-Boc-1,2-diaminoethane (1.2 equiv.) was dissolved in DCM and reacted with NBD-Cl (1.0 equiv.) at room temperature. DIEA was added dropwise to maintain the pH at approximately 8. After stirring for 6 h, the reaction mixture was diluted with DCM, and the organic phase was sequentially washed with aqueous $NaHCO_3$ and water. The organic layer was then air-dried, and the crude product was treated with 80% TFA in DCM for 1 h. Finally, the reaction mixture was evaporated to dryness, redissolved in $H_2O$/acetonitrile, and lyophilized.

Solid phase peptide synthesis

Peptides were synthesized using standard Fmoc solid-phase peptide synthesis on 2-chlorotrityl chloride resin with the corresponding Fmoc-protected amino acids. The 2-Cl resin was first swollen in dry DCM for 30 min, after which the initial Fmoc-protected amino acid was loaded for 4 h. Remaining active sites were capped with a solution of DCM/MeOH/DIEA (17:2:1) for 15 min. The Fmoc group was removed with 20% piperidine in DMF for 20 min, and subsequent Fmoc-protected amino acids were coupled using HBTU as the activating reagent.

After removal of the Fmoc group from the second amino acid (or the third amino acid for peptide **3**), 3-(acetylthio)propionic acid (for peptide **1**), 3-(tert-butoxy)propanoic acid (for peptide **2**), or acetyl anhydride (for peptide **3**) was coupled to the free amino group. After completion of the synthesis, the peptide was cleaved from the resin using TFA/TIPS/$H_2O$ (95:2.5:2.5) for 1 h. The crude peptide solution was air-dried, precipitated with diethyl ether, and centrifuged to obtain the solid product, which was further dried by lyophilization. Lowercase letters denote D-amino acids.

Synthesis of peptides **1-3**

In brief, 0.1 mmol of the synthesized peptide (shown in the dashed box in Scheme S2) was dissolved in 1 mL of dry DMF, and HBTU (0.12 mmol) was added directly to the solution. DIEA was added dropwise to adjust the pH to approximately 8. After stirring for 30 min, NBD-ethylenediamine (0.12 mmol) was added, while maintaining the pH at ~8 with DIEA. The reaction mixture was stirred overnight, after which the solvent was evaporated. The Fmoc group was then removed using 20% piperidine for 20 min. Following a second solvent removal, the crude product was dissolved in MeOH and purified by RP-HPLC to afford the desired compounds.

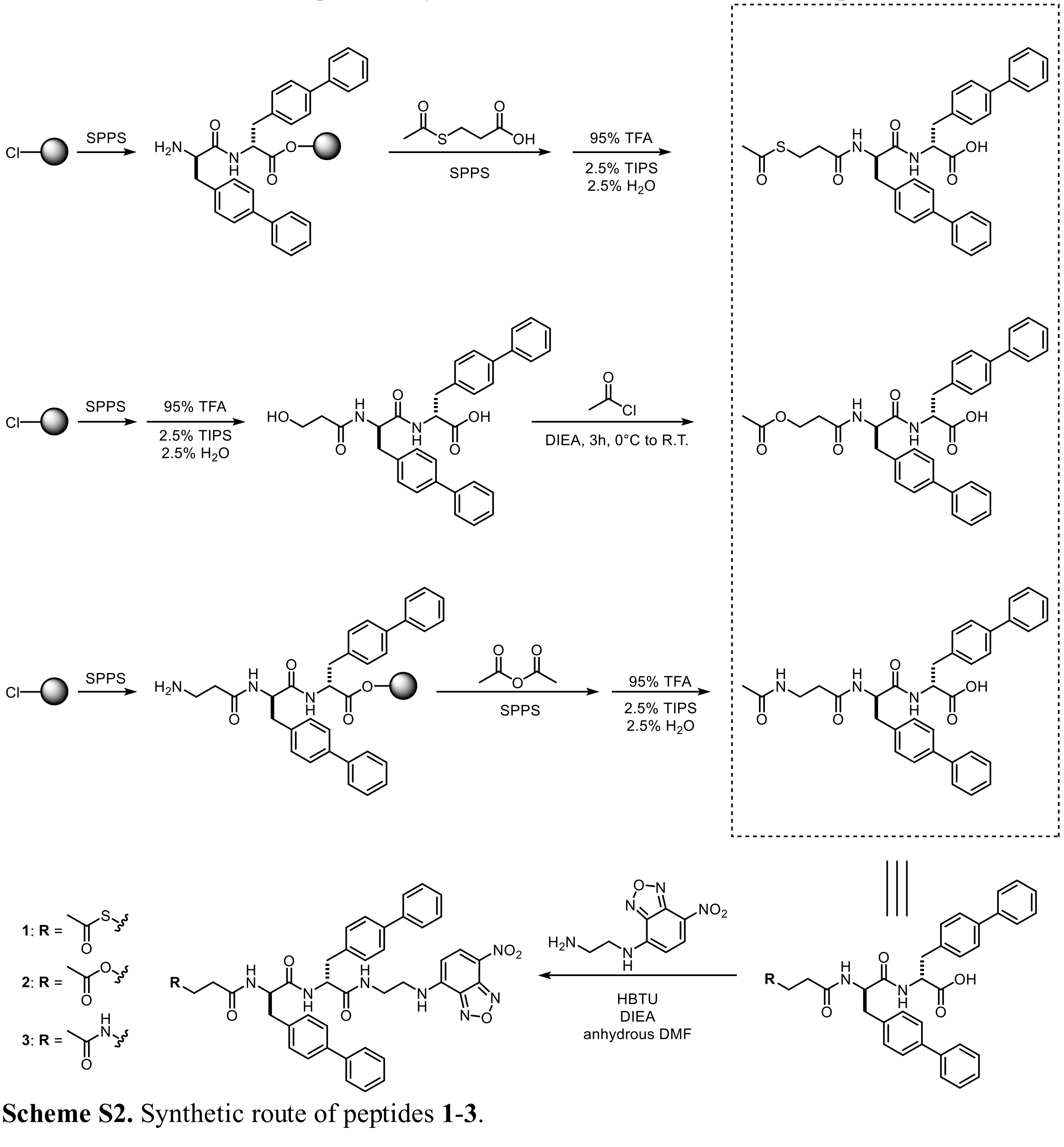


**Scheme S2.** Synthetic route of peptides **1-3**.

Cell culture

HeLa, HepG-2, PANC-1, SH-SY5Y, T98G, H460, A431 and PC-3 cell lines were obtained from the American Type Culture Collection (ATCC; Manassas, VA, USA). HeLa, HepG-2, and T98G cells were cultured in MEM supplemented with 10% fetal bovine serum (FBS) and 1% penicillin-streptomycin (100 U/mL penicillin, 100 μg/mL streptomycin). PANC-1 and A431 cells were cultured in DMEM supplemented with 10% FBS and 1% penicillin–streptomycin. SH-SY5Y cells were cultured in a 1:1 mixture of EMEM and F-12 medium supplemented with 10% FBS and 1% penicillin–streptomycin. H460 and PC-3 cells were cultured in RPMI 1640 medium supplemented with 10% FBS and 1% penicillin–streptomycin. All the cell lines were cultured at 37 °C in a humidified incubator with 5% $CO_2$.

MTT assays

The MTT assay was used to determine cell viability for cytotoxicity evaluation. Cells were seeded in 96-well plates at a density of $1 \times 10^4$ cells per well and allowed to adhere for 24 h. The medium was then replaced with fresh medium containing varying concentrations of compounds. After 24, 48, or 72 h of treatment, the compound-containing medium was removed, and 100 μL of fresh medium supplemented with 10 μL of MTT solution (5 mg/mL; ACROS Organics) was added to each well. Plates were incubated for an additional 4 h at 37 °C. Subsequently, 100 μL of SDS–HCl solution was added to stop the reaction and dissolve the formazan crystals. Absorbance at 595 nm was measured using a DTX880 multimode detector. All experiments were performed in triplicate (n = 3), and cell viability was calculated as a percentage relative to untreated controls.

Confocal laser scanning microscopy (CLSM) imaging

Confocal dishes (35 mm; 20 mm #1.5 glass-bottom well; Cellvis) were used for CLSM sample preparation. For live-cell imaging, exponentially growing cells were seeded at $1.0 \times 10^5$ cells per dish and cultured for 24 h. The medium was then replaced with fresh medium containing the compound of interest, and cells were incubated at 37 °C in a humidified atmosphere with 5% $CO_2$ for the indicated time. Live-cell images were acquired using a Nikon AX-R confocal laser scanning microscope (CLSM).

For time-lapse imaging, exponentially growing cells were seeded on confocal dishes at the same density and cultured for 24 h. Cells were washed three times with Live Cell Imaging Solution, stained with Hoechst 33342 for 10 min, and washed four additional times to remove excess dye. Time-lapse imaging was performed on a Nikon AX-R CLSM. Cell positions and the laser focal plane were set using nuclear fluorescence excited by the 405 nm laser, and the Nikon Perfect Focus System was used to minimize focus drift. The imaging solution was then replaced with fresh imaging solution containing the compound of interest, and multichannel images were acquired immediately. Time-series images were collected at 1 min intervals for 40 cycles, and the fluorescence images were saved for subsequent analysis.

Plasmid transfection

96-well glass-bottom plates (#1.5 cover glass; Cellvis) were used for plasmid transfection. Cells were seeded at $8 \times 10^3$ cells per well and cultured for 24 h to allow attachment. At 40–50% confluency, cells were transfected using Xfect™ Transfection Reagent. Briefly, 3 μg of plasmid DNA was mixed with 65 μL of Xfect Reaction Buffer, followed by addition of 1 μL Xfect Polymer. The mixture was gently vortexed and incubated at room temperature for 10 min to allow nanoparticle complex formation. Then, 10 μL of the transfection complex was added dropwise to each well and the plate was gently rocked to ensure even distribution. Cells were incubated overnight at 37 °C, after which the medium was replaced with fresh culture medium and incubation was continued for an additional 48 h. Cells were then prepared for live-cell imaging.

Drug resistance test

HeLa cells were divided into two groups. Group 1 was treated with compound **1** (200 nM) for 24 h, after which the medium was replaced with fresh medium, and cells were allowed to recover and proliferate for an additional 2 days. Surviving cells were then subcultured and expanded to 80–90% confluency before being re-treated with compound **1** at gradually increasing concentrations (up to 2 μM), for a total of seven additional stimulation cycles. Group 2 was treated with vehicle (DMSO) and subjected to the same procedure as a control. After eight cycles, surviving cells from Group 1 and control cells from Group 2 were harvested and seeded into 96-well plates at $1 \times 10^5$ cells per well. After 24 h, cells were treated with compound **1** for 24 h, and cell viability was determined using the MTT assay.

Untargeted lipidomics profiling

Lipidomics samples were prepared according to a previously reported protocol.[1] Briefly, $1.2 \times 10^7$ HeLa cells were treated with compound **1** (2 μM) or vehicle control for 6 h or 12 h. Cells were harvested at room temperature by trypsinization and centrifugation (1000 × g, 5 min). After removal of the supernatant, the cell pellet was resuspended in 200 μL of 1× PBS.

For extraction of nonpolar lipids, 1.5 mL of HPLC-grade methanol was added and the suspension was vortexed for 1 min, followed by addition of 5 mL of methyl tert-butyl ether (MTBE). Samples were rocked for 1 h at room temperature. Phase separation was induced by addition of 1.2 mL water, followed by vortexing for 1 min and centrifugation (1000 × g, 10 min). The upper MTBE phase was collected, and the lower aqueous phase was re-extracted with two volumes of MTBE/methanol/water (10:3:2.5, v/v/v). The organic phases were combined, dried under a nitrogen stream, and submitted for MS analysis at the BIDMC–Harvard Mass Spectrometry Facility. Lipidomics data were median-normalized within each sample.

FRAP assay

FRAP was performed on a Nikon AX-R confocal laser scanning microscope using a 60×/1.4 NA oil-immersion objective. Cells were treated with compound **1** (2 μM) for 4 h, and cells displaying whorls were selected for analysis. Ten pre-bleach images were acquired, after which a defined region within an individual whorl was photobleached using the 488 nm laser at 100% power. Time-lapse images (512 × 512 pixels) were then collected at 0.54 s intervals with the 488 nm excitation, using a pinhole of 1 Airy unit, and acquisition was continued until the recovery reached a plateau. Fluorescence recovery in the bleached region was background-corrected, normalized to the pre-bleach intensity, and fit to a single-exponential function.

ER-to-Golgi anterograde trafficking analysis

HeLa cells seeded on confocal dishes were transfected with RUSH constructs (Str-KDEL_TNF-SBP-mCherry and Str-KDEL_SBP-mCherry–E-cadherin). After transfection, cells were treated with compound **1** (2 μM) for 4 h at 37 °C and stained with Hoechst 33342. Cells were then transferred to the confocal microscope for live-cell imaging, and the culture medium was replaced with fresh medium containing biotin (40 μM) to trigger synchronized cargo release. Time-lapse imaging was initiated immediately after biotin addition using time-series mode. Images were saved and analyzed offline.

Immunocytochemistry

Culture medium was removed and cells were washed twice with PBS. Cells were fixed with 4% paraformaldehyde for 10 min and permeabilized with 0.1% Triton X-100 in PBS for 7 min. After permeabilization, cells were blocked with 1% BSA and 22.5 mg/mL glycine in PBST for 1 h at room temperature. Primary antibodies were diluted 1:200 in 1% BSA in PBST and incubated with cells overnight at 4 °C. After washing, cells were incubated with Alexa Fluor 647–conjugated secondary antibodies (1:1000 in 1% BSA in PBST) for 1 h at room temperature. Between each step, except after blocking, cells were washed three times with PBS. The cells were then ready for CLSM imaging.

Palmitoylation of CyMA characterized by LC/HR-MS

A total of $1.2 \times 10^7$ cells were treated with compound **1** (2 μM) or vehicle (DMSO) for 2h, then washed twice with HEPES buffer. Cells were harvested and centrifuged to obtain a pellet, which was resuspended in 500 μL HEPES buffer. Dichloromethane (DCM, 1.5 mL) and methanol (1.0 mL) were added, and the mixture was incubated for 10 min at room temperature. Additional DCM (0.5 mL) and Tris–HCl (0.5 mL, 50 mM, pH 2.0) were then added, followed by centrifugation to separate phases. The organic phase was collected, washed with 2.0 mL of a methanol/Tris–HCl mixture (1 mL methanol + 1 mL Tris–HCl, 50 mM, pH 2.0), and centrifuged again to recover the organic layer. The combined organic extract was evaporated under a gentle stream of N2, and the residue was reconstituted in 150 μL methanol for LC/HR-MS analysis. The identity of each compound was confirmed by comparing the experimentally observed high-resolution m/z values with the corresponding calculated exact masses. Assignments were further validated by ensuring that the measured isotopic peak distributions matched the theoretical isotope patterns predicted from the elemental formulas.

Immunoblotting

Cells were cultured to 80–90% confluency and lysed on ice with 500 μL of lysis buffer (supplemented with protease inhibitor cocktail and phosphatase inhibitor cocktails) per 10 cm dish. Lysates were briefly sonicated (10 s) and subjected to three freeze–thaw cycles, then clarified by centrifugation at 12,000 rpm for 10 min at 4 °C. The supernatants were collected, mixed with sample loading buffer, and heated at 95 °C for 8 min to denature proteins. Samples were resolved on precast SDS–PAGE gels (140 V, 40 min) and transferred to PVDF membranes (100 V, 100 min) in an ice bath. Membranes were blocked for 1 h at room temperature and incubated with primary antibodies (1:1000 dilution) overnight at 4 °C. After three washes with TBST (5 min each), membranes were incubated with secondary antibodies (1:20,000 dilution) for 1 h at room temperature. Membranes were then washed with TBST six times, developed with chemiluminescent substrate for 1 min, and imaged using a blot scanner. For reprobing, membranes were stripped with Restore™ Western Blot Stripping Buffer, re-blocked, and incubated with the next primary antibody.

RNA-seq analysis

RNA samples from vehicle control and **1**-treated (2 μM) cells collected at 6 h and 12 h were submitted to Azenta-GENEWIZ for bulk mRNA sequencing. Libraries were prepared using a poly(A)-selection workflow and sequenced on an Illumina HiSeq platform in a paired-end 2 × 150 bp format. Raw reads were processed by trimming adapter sequences and low-quality bases using Trimmomatic (v0.36), and the cleaned reads were aligned to the *Homo sapiens* GRCh38 reference genome from Ensembl using STAR (v2.5.2b). Unique gene hit counts were generated using featureCounts from the Subread package (v1.5.2) based on the gene_id feature in the annotation file. Differential gene expression analysis was performed using DESeq2 with the Wald test. Genes with an adjusted p value $< 0.05$ and an absolute log2 fold change $> 1$ were considered differentially expressed.

Transferrin pulse-chase assay

HeLa cells were treated with compound **1** at the indicated concentrations or vehicle for 4 h, followed by washout. Cells were then incubated with Alexa Fluor 647-labeled transferrin (25 µg/mL) for 1 h, washed, and transferred to fresh medium containing unlabeled transferrin (2.5 mg/mL) for the chase period. Time-lapse CLSM imaging was initiated immediately after the start of the chase, and intracellular AF647 fluorescence was quantified under identical imaging settings.

Filipin III staining

HeLa cells were treated with compound **1** or vehicle for 4 h, fixed with paraformaldehyde for 10 min, and washed three times with PBS. Cells were then stained with Filipin III (50 µg/mL) for 1 h at room temperature in the dark, followed by three washes with PBS. Fluorescence images were acquired by confocal microscopy.

Quantification and data analysis

All graphs were generated using Origin 2021. Fluorescence intensity was quantified using Fiji. Error bars represent the standard deviation unless otherwise noted. Statistical significance between two groups was assessed using two-tailed Student's T-tests. All cell-based and cell-free experiments were independently repeated two to three times.

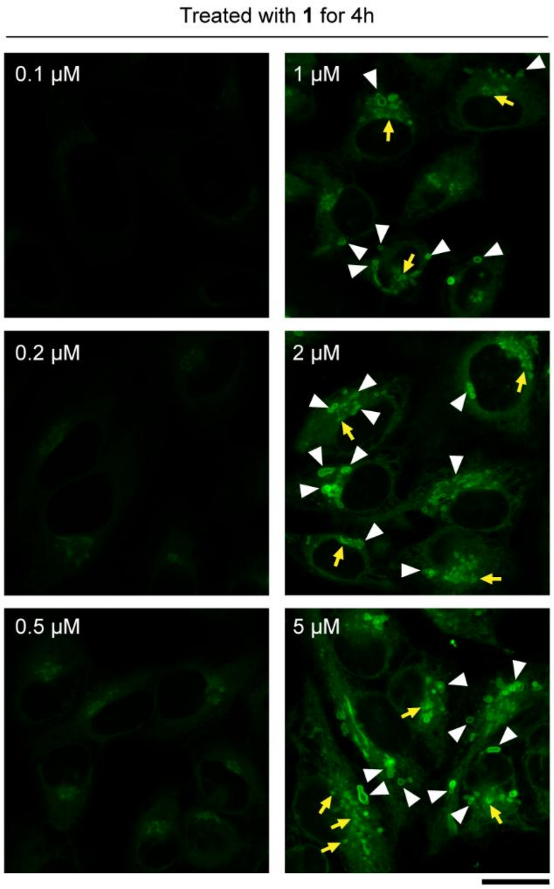


**Fig. S1. Dose- and time-dependent whorl formation in HeLa cells treated with 1.**
Representative CLSM images of HeLa cells treated with gradient concentrations of **1** for 4 h. (Scale bar = 20 μm)

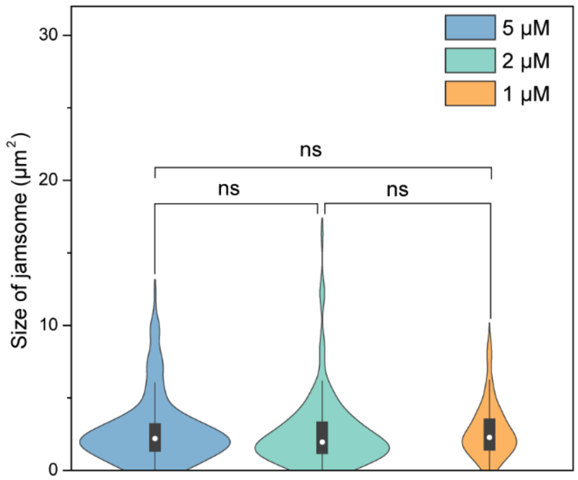


**Fig. S2. Quantification of whorl morphology in HeLa cells treated with 1.**
Quantification of whorl size/morphology distributions in HeLa cells treated with **1** for 4 h.

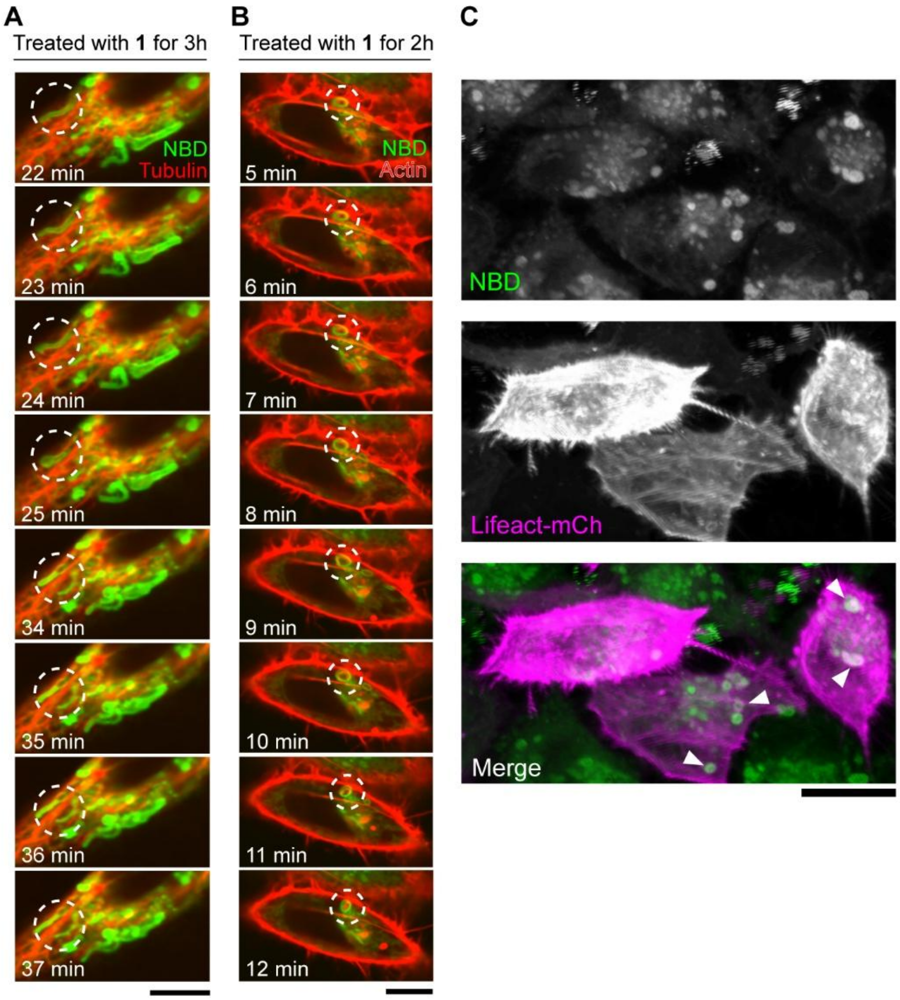


**Fig. S3. Formation of whorls showing transient cytoskeletal interactions.**
(A) Time-lapse CLSM images of HeLa cells stained with Tubulin Tracker Deep Red and treated with **1** (2 μM, 3 h). (Scale bar = 10 μm) (B) Time-lapse CLSM images of HeLa cells stained with CellLight Actin Deep Red and treated with **1** (2 μM, 2 h). (Scale bar = 10 μm) (C) Z-stack CLSM images of HeLa cells transfected with Lifeact-mCh and treated with **1** (5 μM, 4h). (Scale bar = 20 μm) White arrowheads indicate colocalization between whorls and Lifeact-mCh.

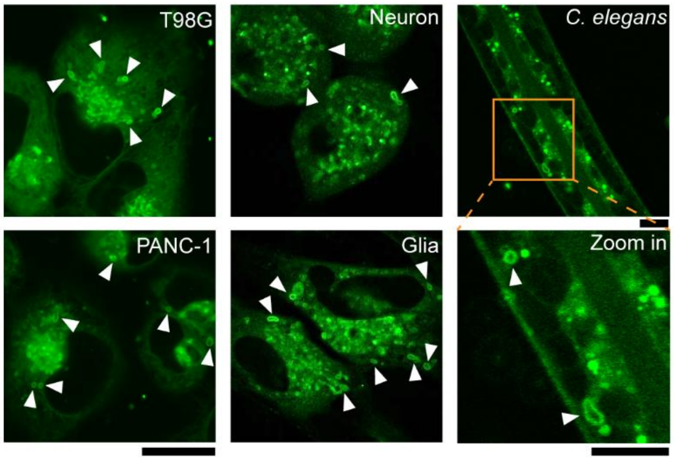


**Fig. S4. Whorl formation in other cancer cell lines, primary mouse neurons, glia, and *C. elegans*.**
Representative images showing whorl formation in other cancer cell lines, primary mouse neurons and glial cells, and in *C. elegans*, following treatment with **1**. (Scale bar = 20 μm)

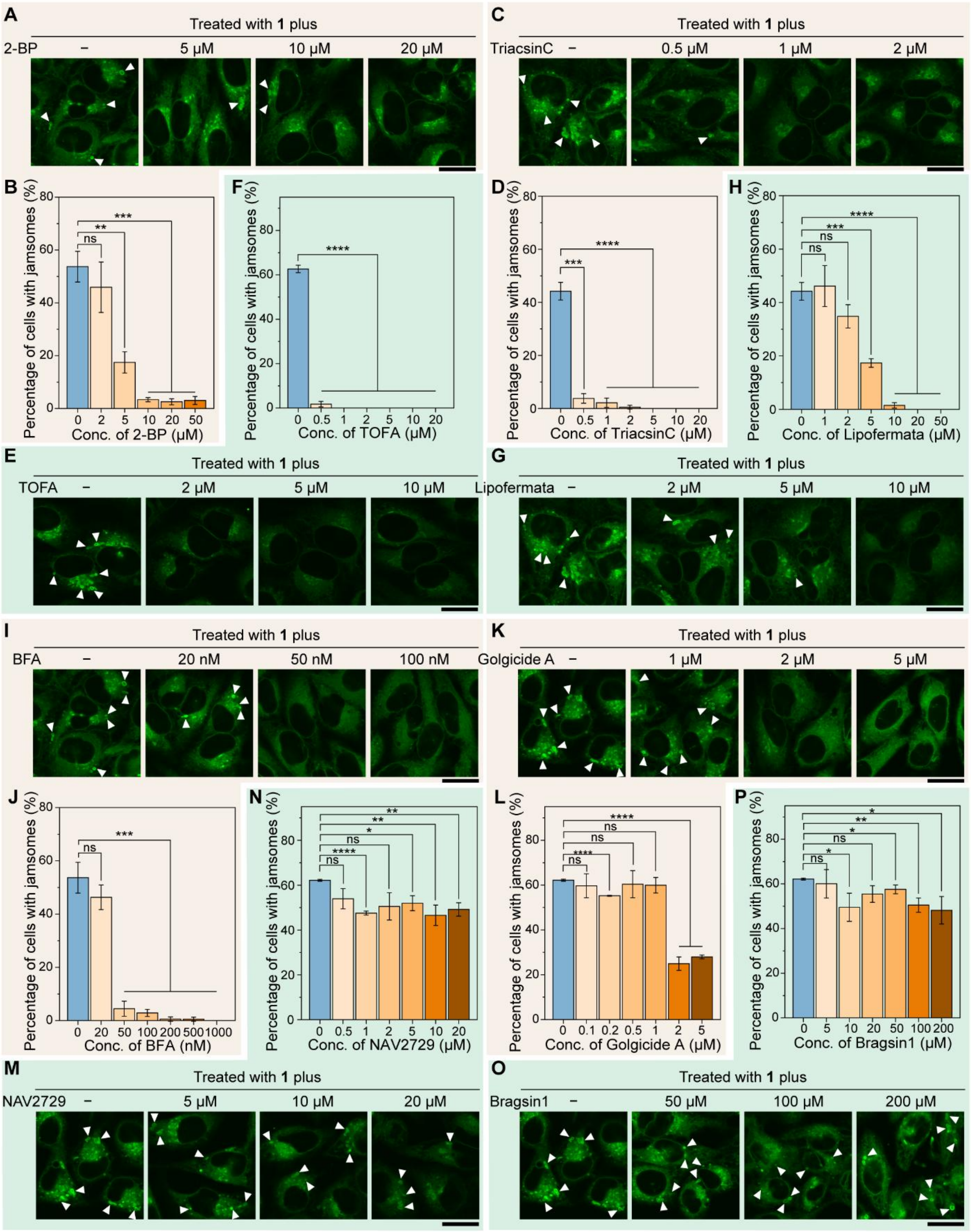


**Fig. S5. Effects of lipid metabolism and Golgi trafficking inhibitors on whorl formation.** (A, C, E, G, I, K, M, O) CLSM images and (B, D, F, H, J, L, N, P) quantification of whorls in cells pretreated with 2-BP (A, B), Triacsin C (C, D), TOFA (E, F), lipofermata (G, H), BFA (I, J), golgicide A (K, L), NAV-2729 (M, N), or bragsin1 (O, P) for 30 min, followed by co-treatment with **1** (2 μM) and the corresponding inhibitor for 4 h. (Scale bar = 20 μm) White arrowheads indicate whorls.

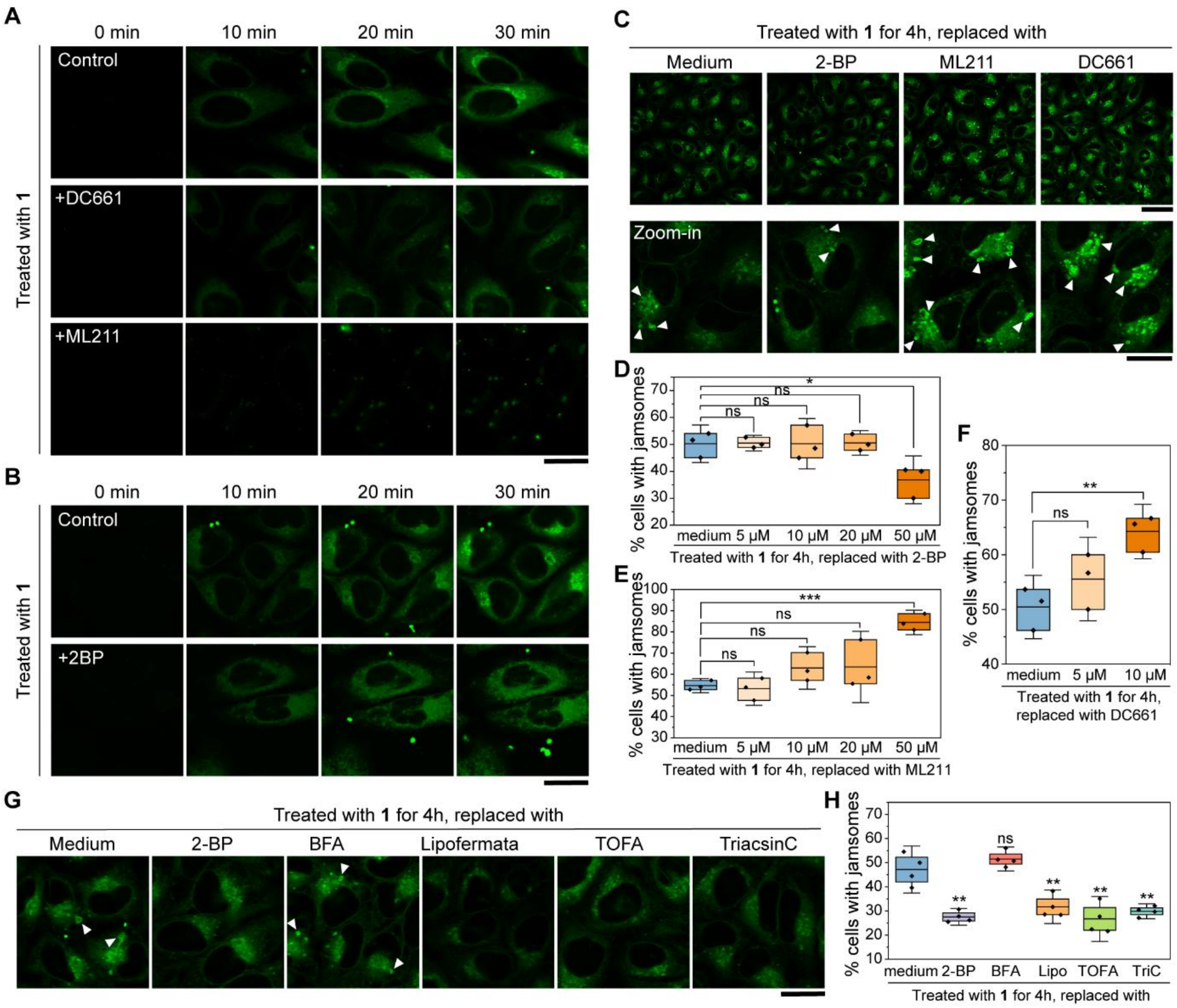


**Fig. S6. Effects of inhibitor treatments on whorl formation.**

(A) Time-lapse CLSM images of HeLa cells treated with **1** (10 μM), with or without pretreatment with DC661 (20 μM, 30 min) or ML211 (50 μM, 30 min). (Scale bar = 20 μm) (B) Time-lapse CLSM images of HeLa cells treated with **1** (10 μM), with or without pretreatment with 2-BP (10 μM, 30 min). (Scale bar = 20 μm) (C) CLSM images and (D–F) quantification of whorls in HeLa cells treated with **1** (2 μM, 4 h), followed by washout and treatment with the indicated inhibitors for 1 h. (Scale bar = 50 μm for upper panel, 20 μm for lower panel) White arrowheads indicate whorls.

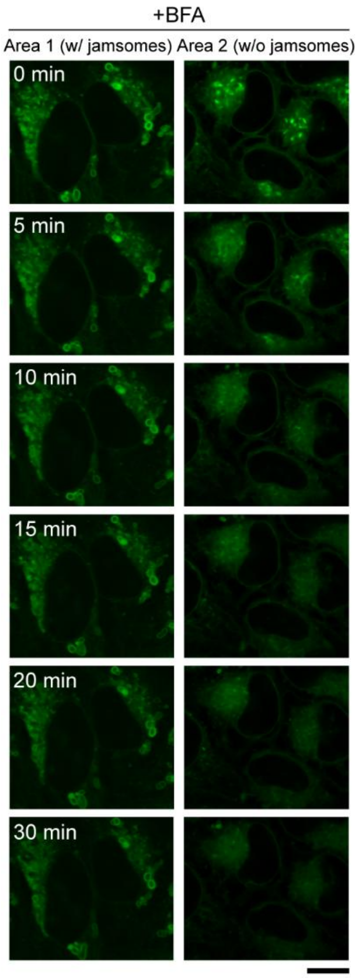


**Fig. S7. Persistence of whorls after washout and BFA treatment.**
Time-lapse CLSM images of HeLa cells treated with **1**, washed out, and then incubated with BFA. (Scale bar = 10 μm)

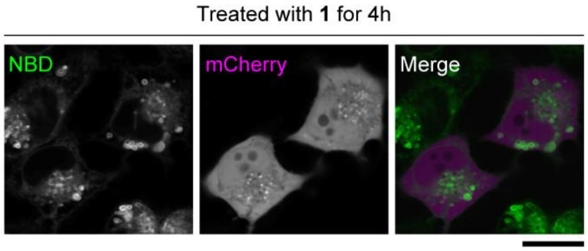


**Fig. S8. A fluorescent-protein-only control does not accumulate on whorls.**
CLSM images of HeLa cells expressing the fluorescent protein alone (no GTPase fusion) and treated with **1**, showing no whorl accumulation. (Scale bar = 20 μm)

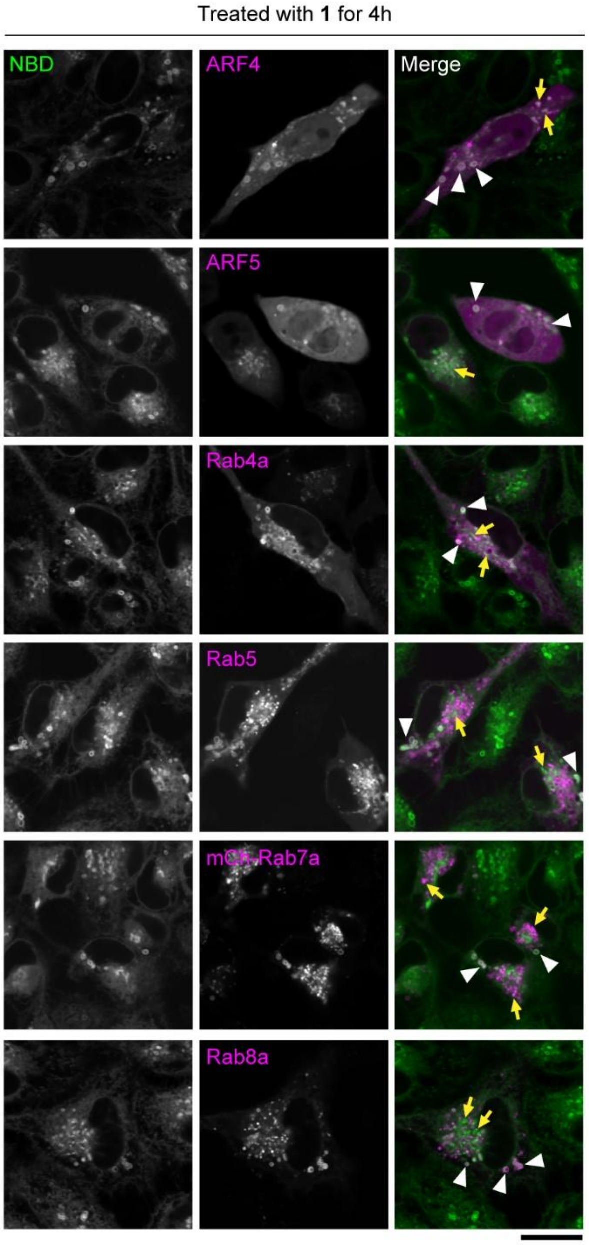


**Fig. S9. Small GTPase recruitment to whorls, extended panel.**
CLSM images of HeLa cells transiently transfected with RFP-tagged small GTPases (ARF and Rab family constructs) and treated with **1** (1 µM, 4 h), extending the set shown in Fig. 2A, B. (Scale bar = 20 µm) White arrowheads indicate whorls; yellow arrows indicate NBD-positive puncta.

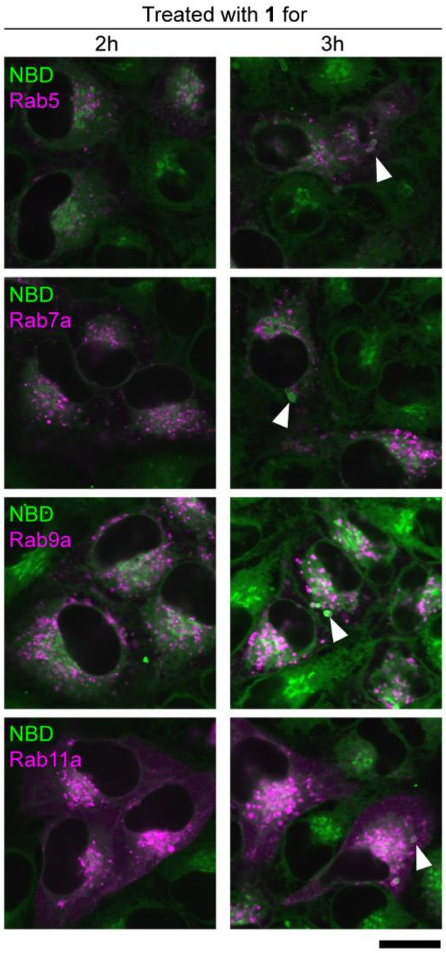


**Fig. S10. Coordinated sequestration of small GTPases on whorls.**
CLSM images of HeLa cells expressing mCh-Rab5, mCh-Rab7a, mCh-Rab9a, or mCh-Rab11a after treatment with **1** (2 μM, 4 h). (Scale bar = 20 μm) White arrowheads indicate whorls.

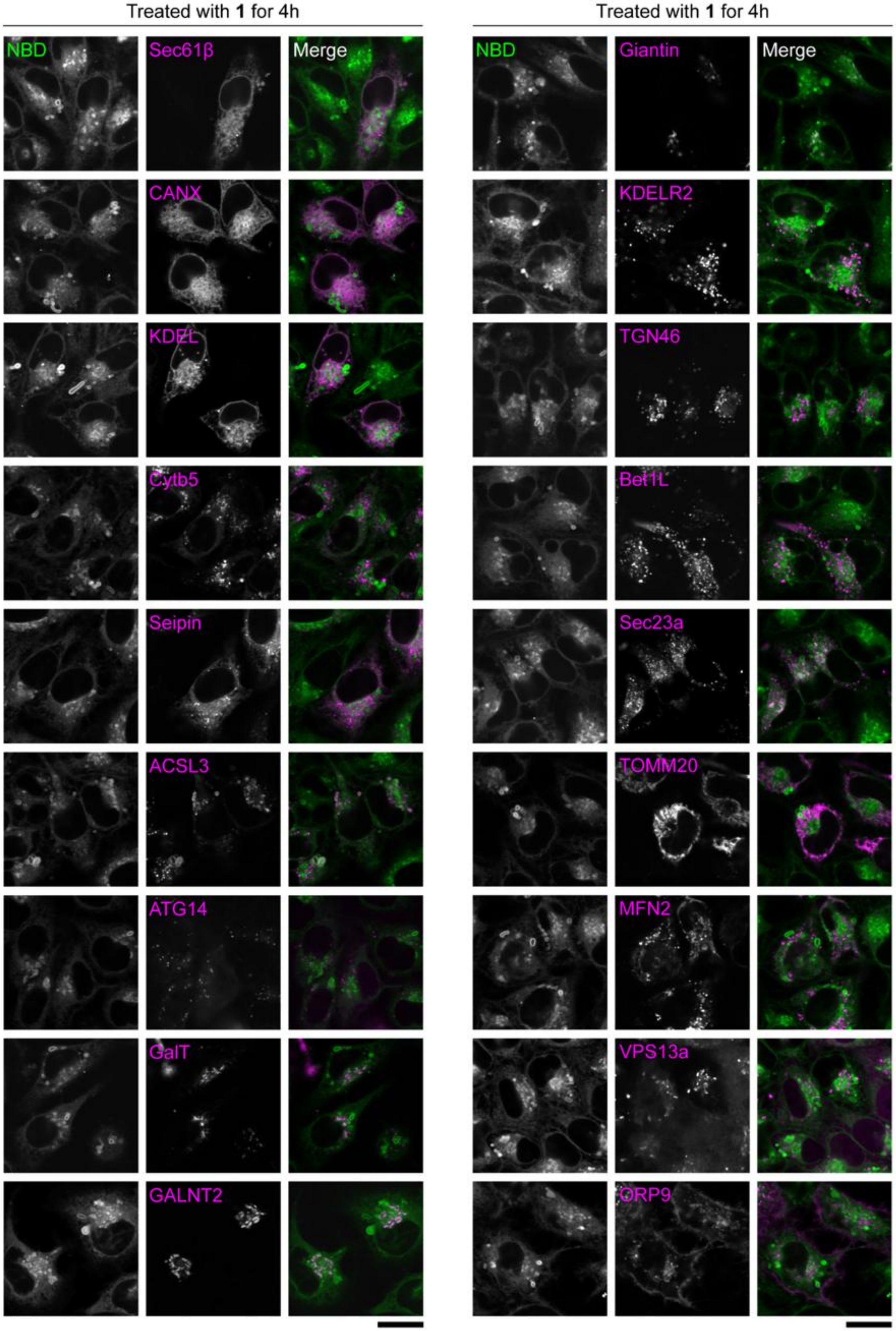

Treated with 1 for 4h
NBD
Sec61β
Merge
CANX
KDEL
Cytb5
Seipin
ACSL3
ATG14
GalT
GALNT2
Treated with 1 for 4h
NBD
Giantin
Merge
KDELR2
TGN46
Bet1L
Sec23a
TOMM20
MFN2
VPS13a
ORP9

**Fig. S11. Non-GTPase proteins (except ACSL3) do not localize to whorls in HeLa cells treated with 1 (Part I).**
CLSM images of HeLa cells transiently transfected with RFP-tagged non-GTPase proteins and treated with **1** (1 µM, 4h). (Scale bar = 20 µm)

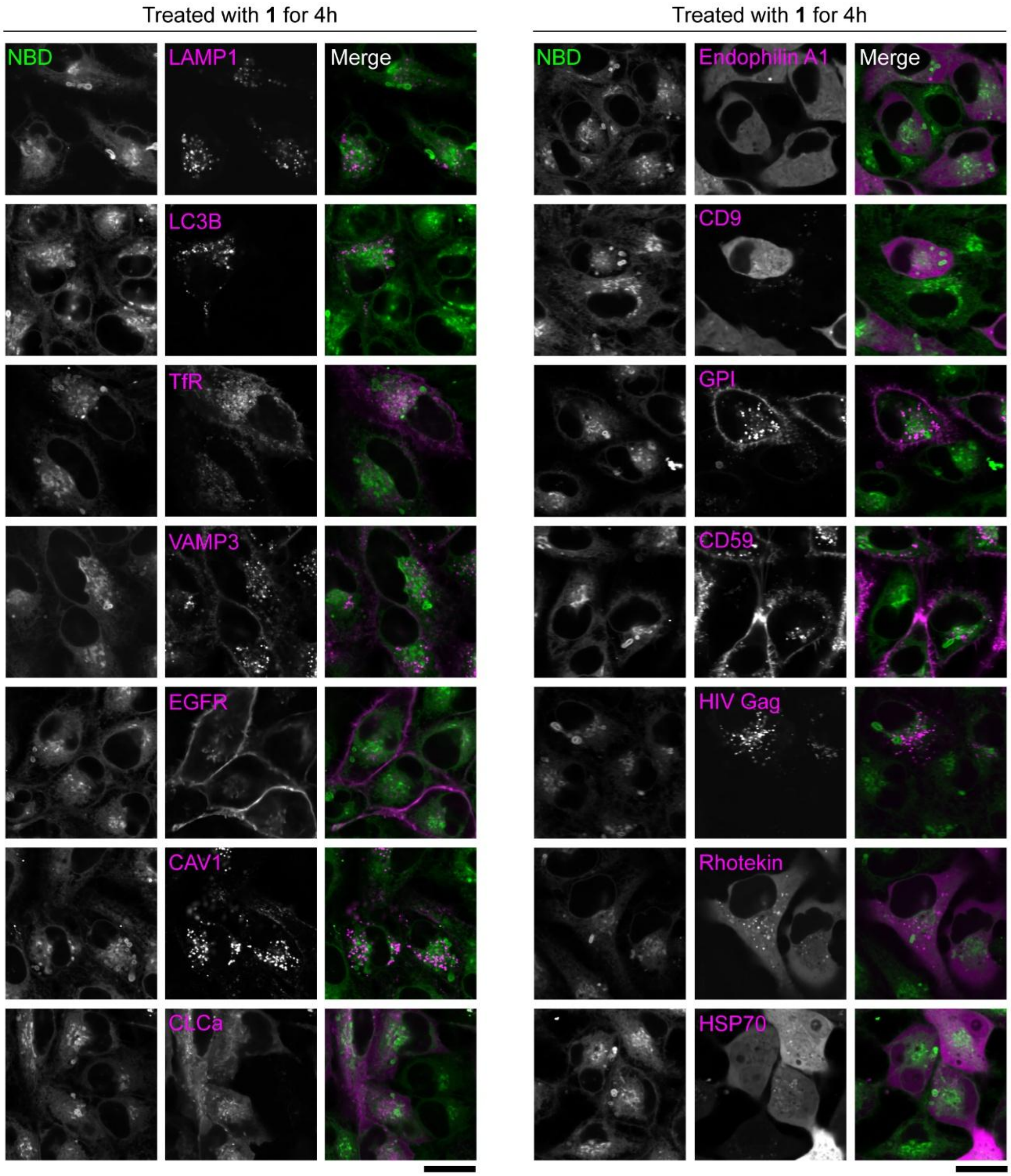


**Fig. S12. Non-GTPase proteins do not localize to whorls in HeLa cells treated with 1 (Part II).**

CLSM images of HeLa cells transiently transfected with RFP-tagged non-GTPase proteins and treated with **1** (1 µM, 4h). (Scale bar = 20 µm)

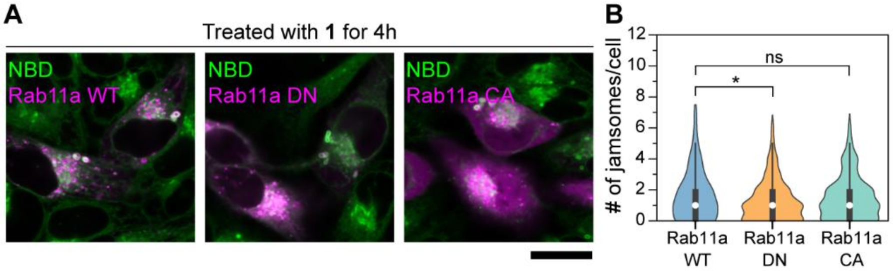


**Fig. S13. Activity-independent sequestration of small GTPases on whorls.**
(A) CLSM images and (B) quantification of whorls in HeLa cells transfected with mCh-Rab11a (WT, S25N dominant-negative mutant, or Q70L constitutively active mutant) and treated with **1** (2 μM, 4 h). (Scale bar = 20 μm)

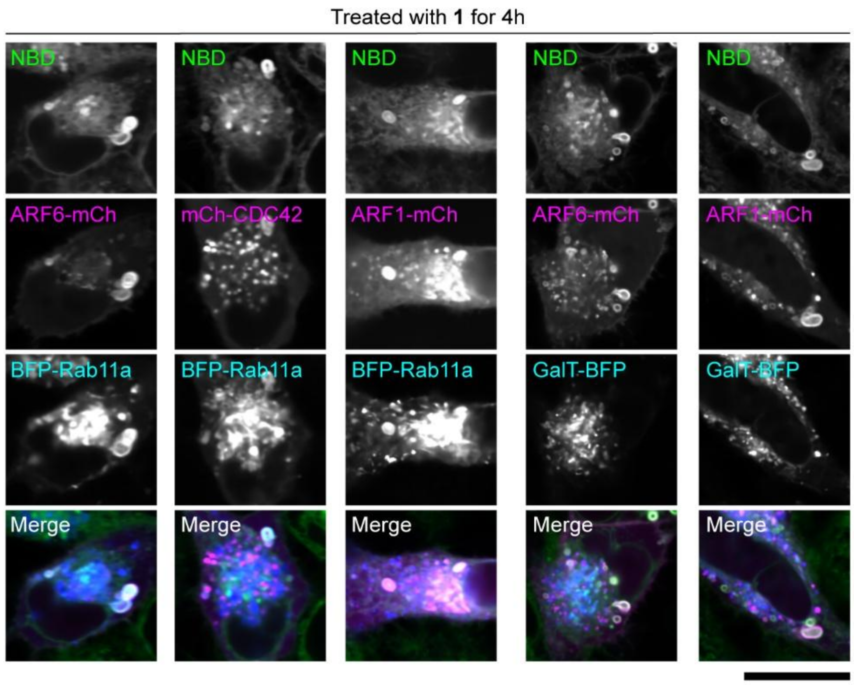


**Fig. S14. Additional co-expression pairs of small GTPases on whorls.**
CLSM images of HeLa cells co-expressing pairs of fluorescently labeled small GTPases or small GTPases together with GalT-BFP after treatment with 1 (2 μM, 4 h). (Scale bar = 20 μm)

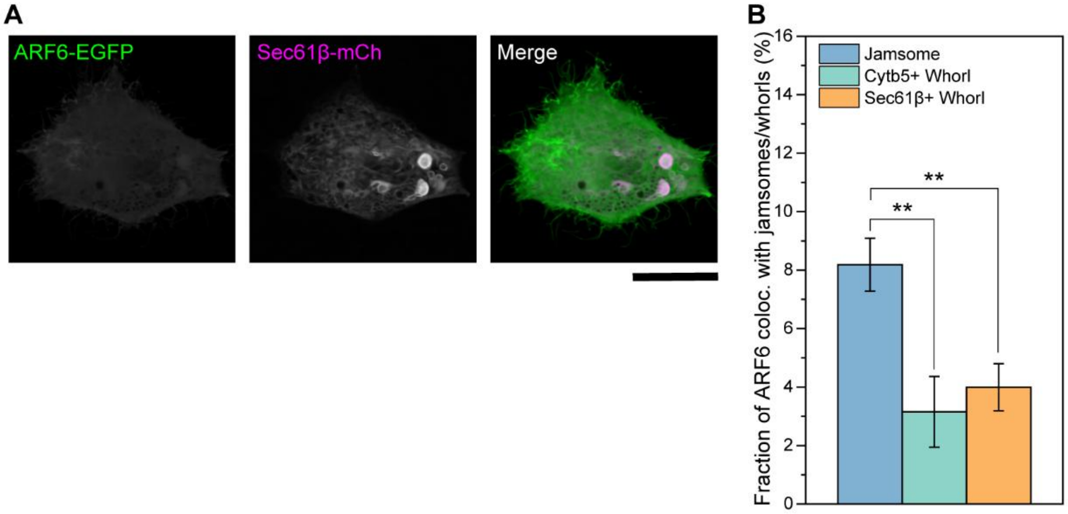


**Fig. S15. ER whorls induced by thapsigargin fail to recruit ARF6.**
(A) CLSM images of ER whorls induced by thapsigargin treatment (2 μM, 6h) and (B) quantification of ER whorls. (Scale bar = 20 μm)

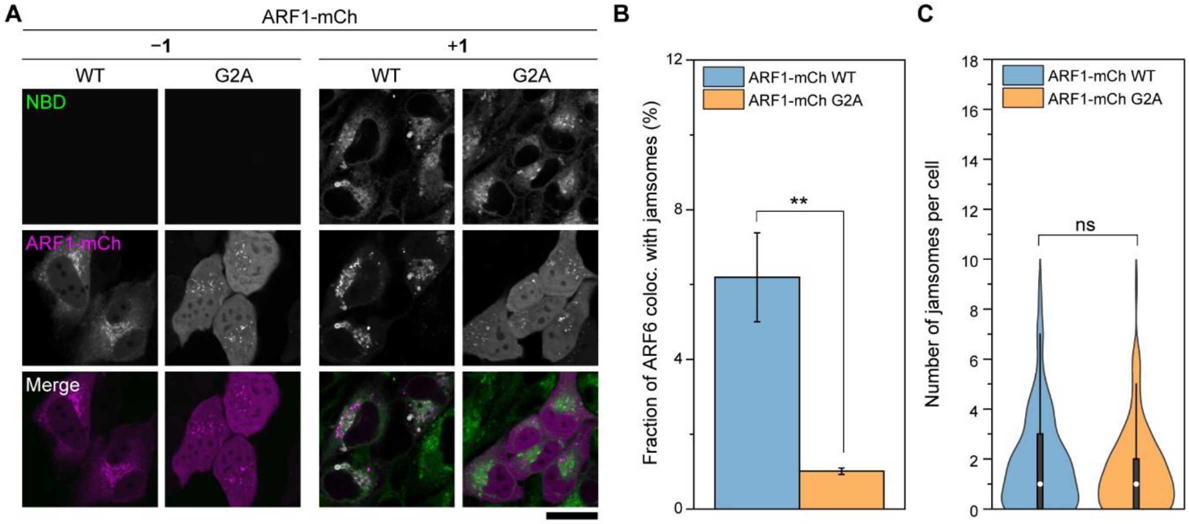


**Fig. S16. An ARF1 G2A mutant accumulates poorly on whorls.**
(A) CLSM images and (B, C) quantification of HeLa cells transiently transfected with ARF1-mCh (WT or G2A) and treated with **1** (2 μM, 4 h). (Scale bar = 20 μm)

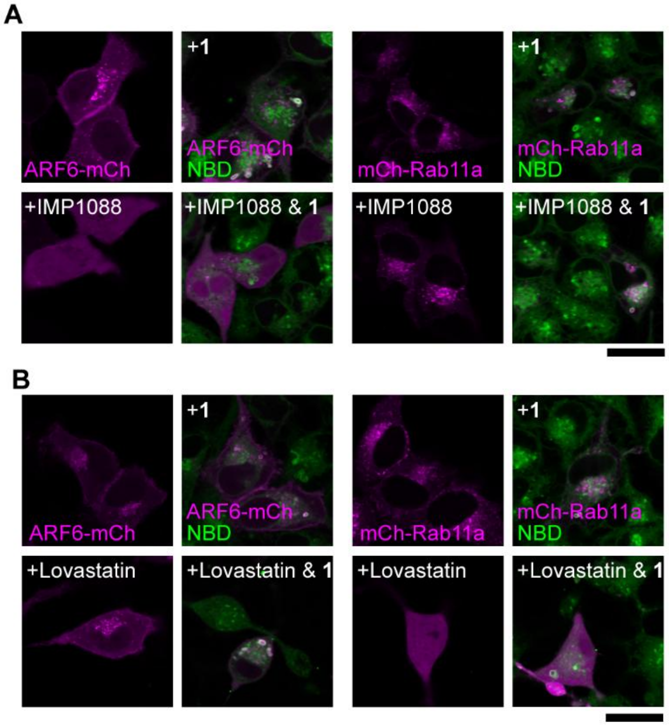


**Fig. S17. Effects of lipidation-related perturbations on whorls biogenesis**
(A) CLSM images of HeLa cells transfected with ARF6-mCh (left) or mCh-Rab11a (right), pretreated with IMP-1088 (20 μM, 24 h), followed by treatment with **1** (2 μM, 4 h). (Scale bar = 20 μm) (B) CLSM images of HeLa cells transfected with ARF6-mCh (left) or mCh-Rab11a (right), pretreated with lovastatin (20 μM, 24 h), followed by treatment with **1** (2 μM, 4 h). (Scale bar = 20 μm)

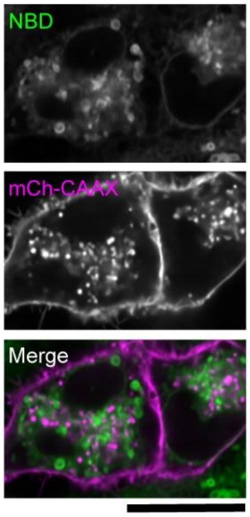


**Fig. S18. A generic prenylated (CAAX) construct does not accumulate on whorls.** CLSM images of HeLa cells transfected with mCh-CAAX and treated with different concentrations of 1 for 4 h. (Scale bar = 20 μm)

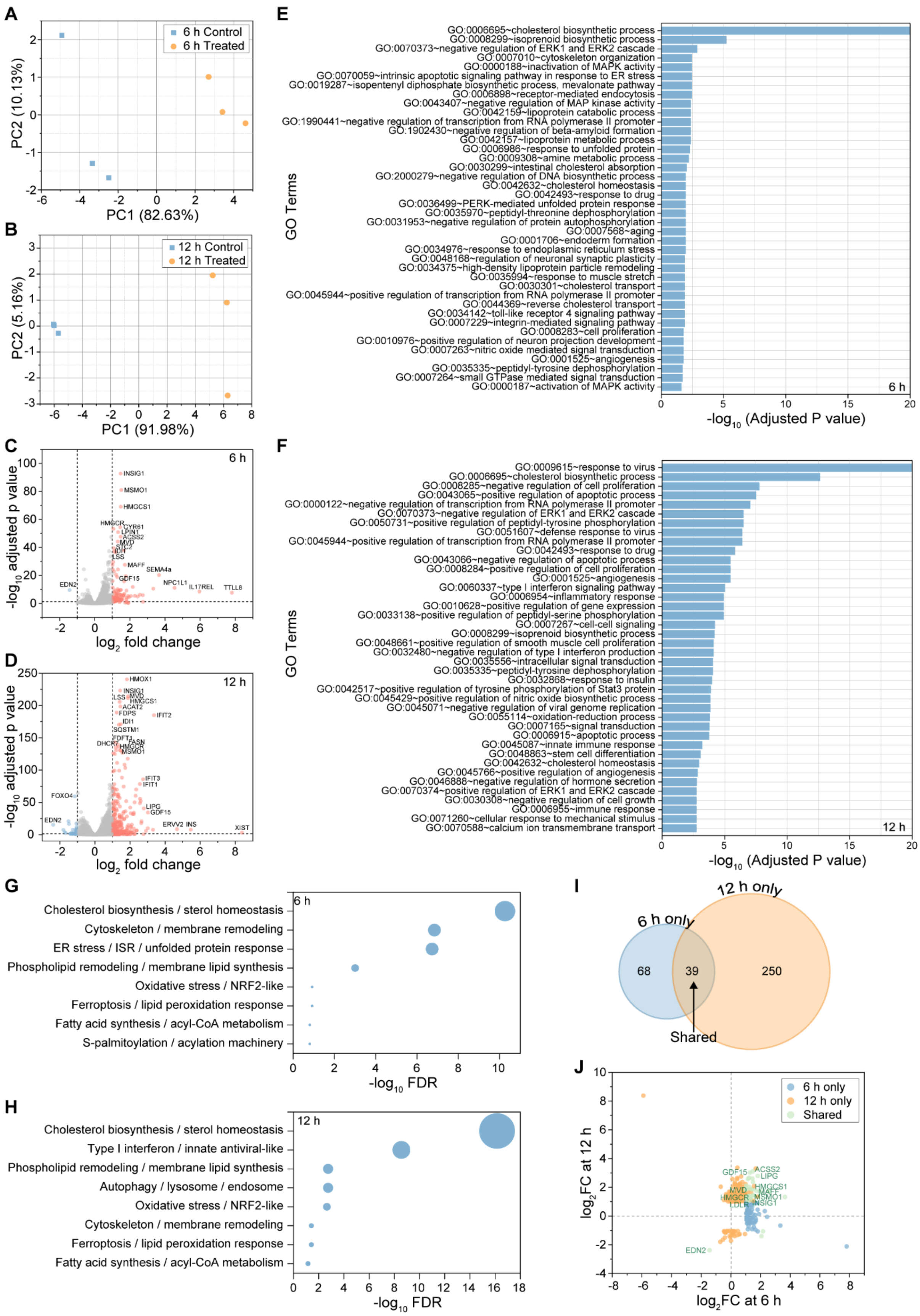
A
6 h Control
6 h Treated
PC2 (10.13%)
PC1 (82.63%)
B
12 h Control
12 h Treated
PC2 (5.16%)
PC1 (91.98%)
C
6 h
-log10 adjusted p value
log2 fold change
INSIG1
MSMO1
HMGCS1
HMGCR
CYR61
LPIN1
ACSS2
MVD
LSS
MAFF
SEMA4a
GDF15
NPC1L1
IL17REL
TTLL8
EDN2
D
12 h
HMOX1
INSIG1
LSS
MVD
HMGCS1
ACAT2
FDPS
IFIT2
IDI1
SQSTM1
FDFT1
FASN
DHCR7
HMGCR
MSMO1
IFIT3
IFIT1
FOXO4
LIPG
GDF15
EDN2
ERVV2
INS
XIST
E
GO Terms
GO:0006695~cholesterol biosynthetic process
GO:0008299~isoprenoid biosynthetic process
GO:0070373~negative regulation of ERK1 and ERK2 cascade
GO:0007010~cytoskeleton organization
GO:0000188~inactivation of MAPK activity
GO:0070059~intrinsic apoptotic signaling pathway in response to ER stress
GO:0019287~isopentenyl diphosphate biosynthetic process, mevalonate pathway
GO:0006898~receptor-mediated endocytosis
GO:0043407~negative regulation of MAP kinase activity
GO:0042159~lipoprotein catabolic process
GO:1990441~negative regulation of transcription from RNA polymerase II promoter
GO:1902430~negative regulation of beta-amyloid formation
GO:0042157~lipoprotein metabolic process
GO:0006986~response to unfolded protein
GO:0009308~amine metabolic process
GO:0030299~intestinal cholesterol absorption
GO:2000279~negative regulation of DNA biosynthetic process
GO:0042632~cholesterol homeostasis
GO:0042493~response to drug
GO:0036499~PERK-mediated unfolded protein response
GO:0035970~peptidyl-threonine dephosphorylation
GO:0031953~negative regulation of protein autophosphorylation
GO:0007568~aging
GO:0001706~endoderm formation
GO:0034976~response to endoplasmic reticulum stress
GO:0048168~regulation of neuronal synaptic plasticity
GO:0034375~high-density lipoprotein particle remodeling
GO:0035994~response to muscle stretch
GO:0030301~cholesterol transport
GO:0045944~positive regulation of transcription from RNA polymerase II promoter
GO:0044369~reverse cholesterol transport
GO:0034142~toll-like receptor 4 signaling pathway
GO:0007229~integrin-mediated signaling pathway
GO:0008283~cell proliferation
GO:0010976~positive regulation of neuron projection development
GO:0007263~nitric oxide mediated signal transduction
GO:0001525~angiogenesis
GO:0035335~peptidyl-tyrosine dephosphorylation
GO:0007264~small GTPase mediated signal transduction
GO:0000187~activation of MAPK activity
6 h
-log10 (Adjusted P value)
F
GO Terms
GO:0009615~response to virus
GO:0006695~cholesterol biosynthetic process
GO:0008285~negative regulation of cell proliferation
GO:0043065~positive regulation of apoptotic process
GO:0000122~negative regulation of transcription from RNA polymerase II promoter
GO:0070373~negative regulation of ERK1 and ERK2 cascade
GO:0050731~positive regulation of peptidyl-tyrosine phosphorylation
GO:0051607~defense response to virus
GO:0045944~positive regulation of transcription from RNA polymerase II promoter
GO:0042493~response to drug
GO:0043066~negative regulation of apoptotic process
GO:0008284~positive regulation of cell proliferation
GO:0001525~angiogenesis
GO:0060337~type I interferon signaling pathway
GO:0006954~inflammatory response
GO:0010628~positive regulation of gene expression
GO:0033138~positive regulation of peptidyl-serine phosphorylation
GO:0007267~cell-cell signaling
GO:0008299~isoprenoid biosynthetic process
GO:0048661~positive regulation of smooth muscle cell proliferation
GO:0032480~negative regulation of type I interferon production
GO:0035556~intracellular signal transduction
GO:0035335~peptidyl-tyrosine dephosphorylation
GO:0032868~response to insulin
GO:0042517~positive regulation of tyrosine phosphorylation of Stat3 protein
GO:0045429~positive regulation of nitric oxide biosynthetic process
GO:0045071~negative regulation of viral genome replication
GO:0055114~oxidation-reduction process
GO:0007165~signal transduction
GO:0006915~apoptotic process
GO:0045087~innate immune response
GO:0048863~stem cell differentiation
GO:0042632~cholesterol homeostasis
GO:0045766~positive regulation of angiogenesis
GO:0046888~negative regulation of hormone secretion
GO:0070374~positive regulation of ERK1 and ERK2 cascade
GO:0030308~negative regulation of cell growth
GO:0006955~immune response
GO:0071260~cellular response to mechanical stimulus
GO:0070588~calcium ion transmembrane transport
12 h
-log10 (Adjusted P value)
G
6 h
Cholesterol biosynthesis / sterol homeostasis
Cytoskeleton / membrane remodeling
ER stress / ISR / unfolded protein response
Phospholipid remodeling / membrane lipid synthesis
Oxidative stress / NRF2-like
Ferroptosis / lipid peroxidation response
Fatty acid synthesis / acyl-CoA metabolism
S-palmitoylation / acylation machinery
-log10 FDR
H
12 h
Cholesterol biosynthesis / sterol homeostasis
Type I interferon / innate antiviral-like
Phospholipid remodeling / membrane lipid synthesis
Autophagy / lysosome / endosome
Oxidative stress / NRF2-like
Cytoskeleton / membrane remodeling
Ferroptosis / lipid peroxidation response
Fatty acid synthesis / acyl-CoA metabolism
-log10 FDR
I
12 h only
6 h only
68
39
250
Shared
J
6 h only
12 h only
Shared
log2FC at 12 h
log2FC at 6 h
GDF15
ACSS2
LIPG
MVD
HMGCS1
HMGCR
MAFF
MSMO1
LDLR
INSIG1
EDN2

**Fig. S19. RNA-seq analysis of cells treated with 1 for 6 h and 12 h.**

Principal component analysis of transcriptomes at (A) 6 h and (B) 12 h. Volcano plots of differentially expressed genes at (C) 6 h and (D) 12 h. Gene Ontology enrichment analysis of differentially expressed genes at (E) 6 h and (F) 12 h. Curated pathway analysis of biological processes related to lipid homeostasis, membrane remodeling, and stress responses at (G) 6 h and (H) 12 h. (I) Overlap of differentially expressed genes between 6 h and 12 h. (J) Comparison of $\log_2$ fold changes of union differentially expressed genes at 6 h and 12 h.

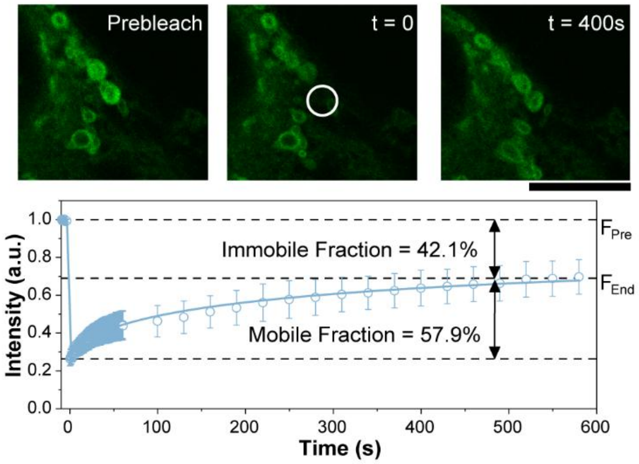


**Fig. S20. FRAP quantification of NBD-labeled 1 within whorl membranes.**
Fluorescence recovery after photobleaching (FRAP) images and quantification, showing an immobile NBD fraction of approximately 42%.

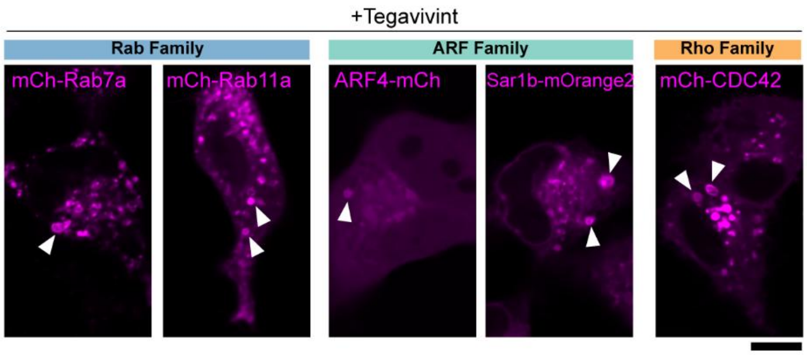


**Fig. S21. Tegavivint recruits additional small GTPases to whorls.**
CLSM images of HeLa cells transfected with RFP-tagged small GTPases (Rab family, ARF family, Rho family) and treated with tegavivint (5 µM, 4 h). (Scale bar = 10 µm) White arrowheads indicate whorls.

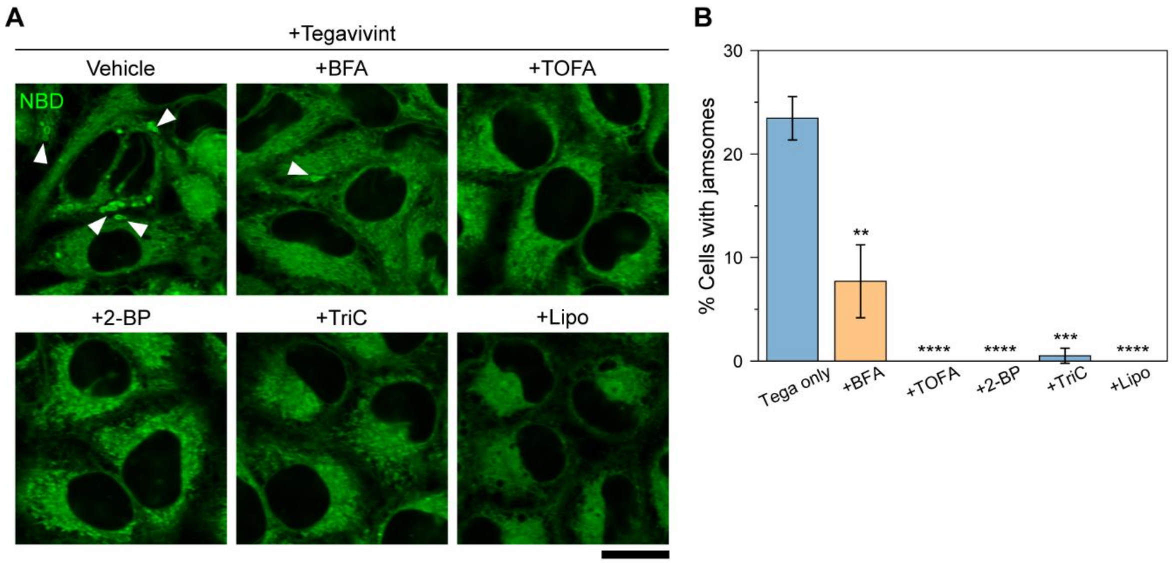


**Fig. S22. Tegavivint-induced whorl formation depends on Golgi integrity, fatty-acid metabolism, and S-acylation.**

CLSM images and quantification of whorls in HeLa cells treated with tegavivint (5 µM, 4 h) in the absence or presence of BFA, TOFA, 2-BP, triacsin C, or lipofermata. (Scale bar = 10 µm) White arrowheads indicate whorls.

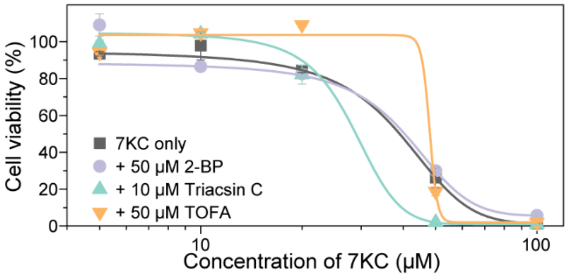


**Fig. S23. Lipid-metabolism inhibitors do not rescue 7-ketocholesterol-induced loss of viability.**
Cell viability of HeLa cells treated with 7-ketocholesterol in the absence or presence of 2-BP, TOFA, or triacsin C for 24 h.

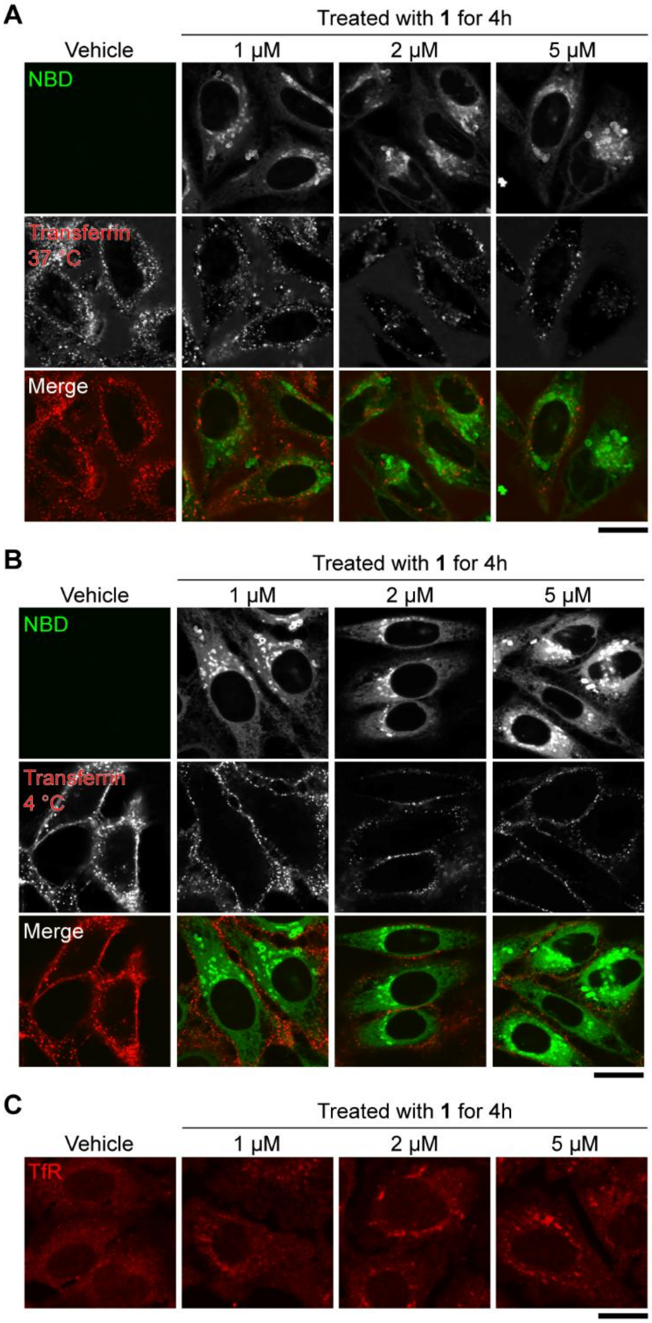


**Fig. S24. Redistribution of the transferrin receptor in HeLa cells treated with 1.**
(A, B) CLSM images of HeLa cells treated with **1** and stained with transferrin AF647. (C) Immunostaining of transferrin receptor in HeLa cells treated with or without **1**. (Scale bar = 20 μm)

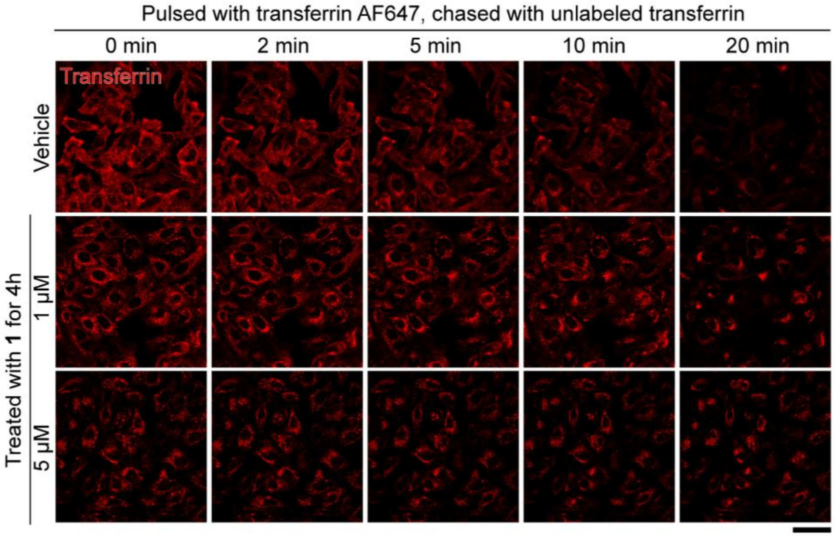


**Fig. S25. Time-lapse imaging supporting the transferrin pulse-chase recycling assay.** Time-lapse CLSM images of HeLa cells treated with **1** in a transferrin pulse-chase assay. (Scale bar = 50 μm)

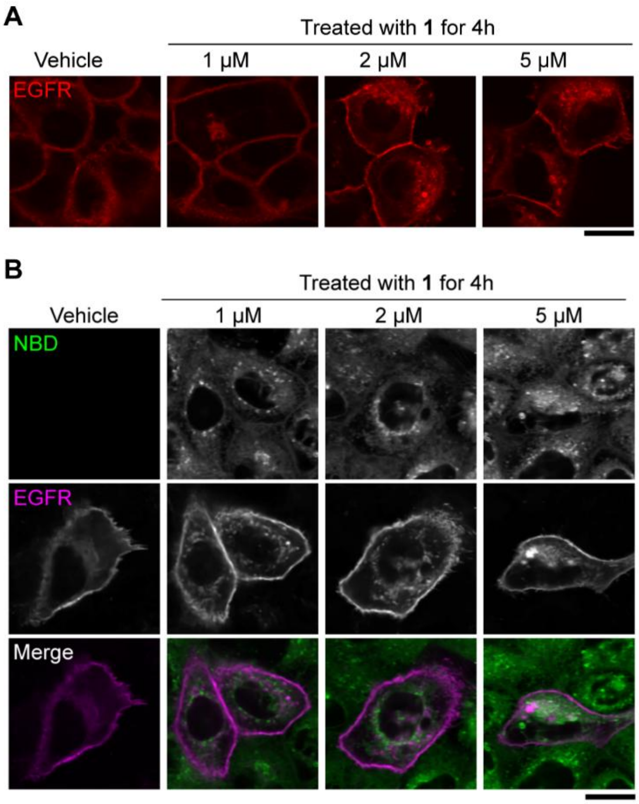


**Fig. S26. Redistribution of EGFR in A431 cells treated with 1.**
(A) Immunostaining of EGFR in A431 cells treated with or without **1**. (B) CLSM images of A431 cells transfected with EGFR-mApple and treated with **1**. (Scale bar = 20 μm)

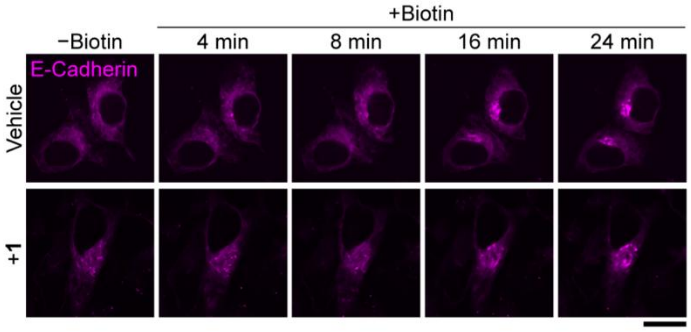


**Fig. S27. Delayed anterograde transport of E-cadherin-mCherry-RUSH in whorl-containing cells.**

Time-lapse CLSM images of HeLa cells expressing E-cadherin-mCherry-RUSH and treated with vehicle or **1** (2 μM, 4 h). (Scale bar = 20 μm)

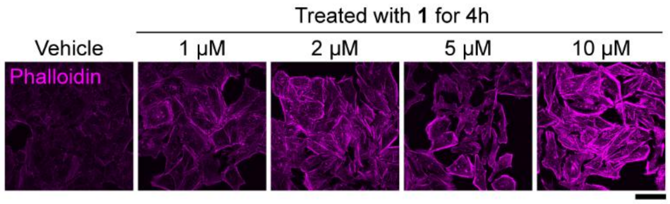


**Fig. S28. Phalloidin staining reveals increased F-actin in cells treated with 1.**
CLSM images of phalloidin AF647 staining in HeLa cells treated with **1**, showing increased F-actin and thickened stress fibers relative to vehicle. (Scale bar = 50 μm)

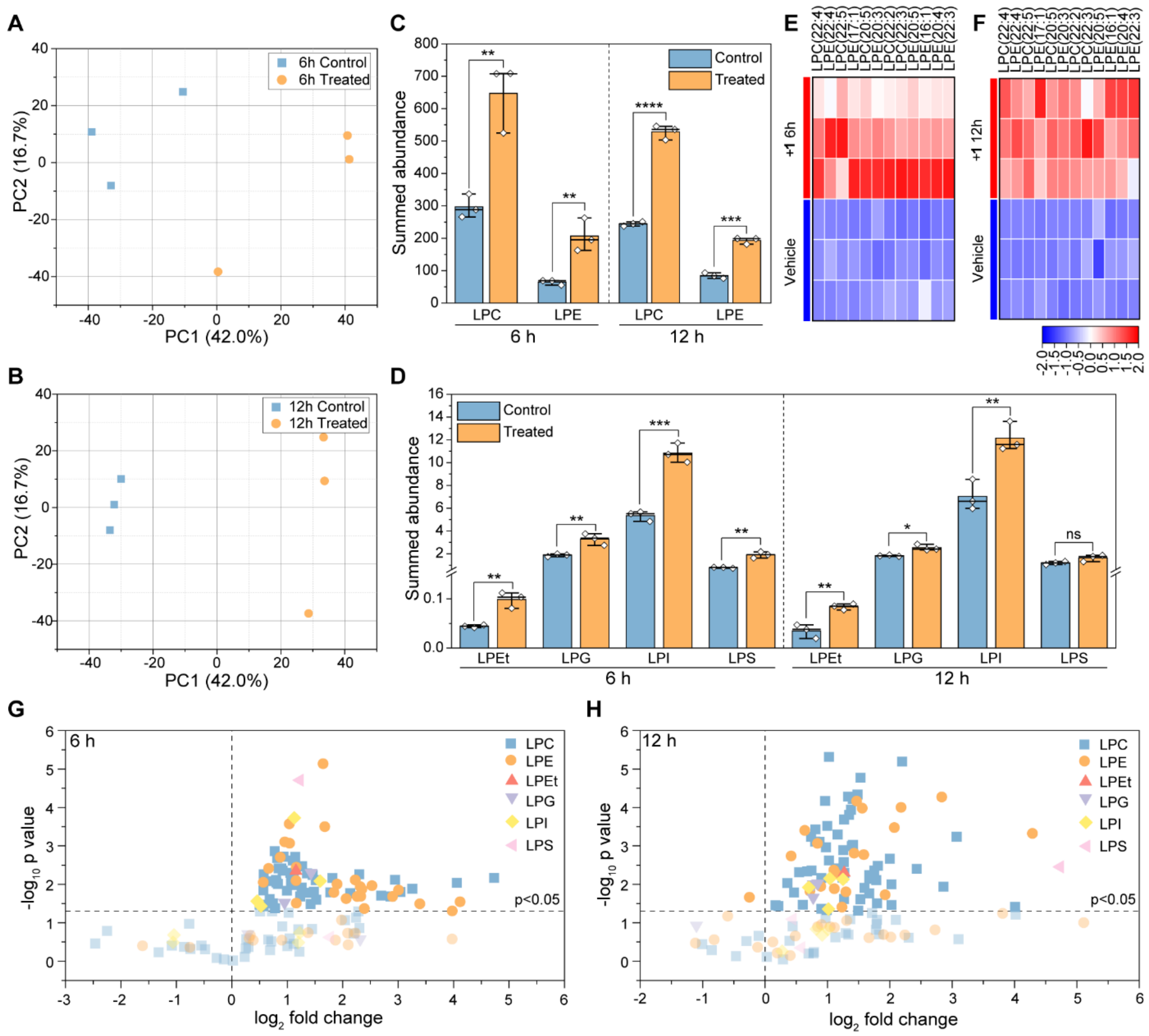

**Fig. S29. Untargeted lipidomics analysis of cells treated with 1 (2 µM).**
Principal component analysis of lipidomes at (A) 6 h and (B) 12 h. (C) Comparison of summed LPC and LPE levels between control and treated groups. (D) Comparison of summed levels of other lysolipid classes between control and treated groups. Top significantly changed lysolipid species at (E) 6 h and (F) 12 h. Volcano plots of all lysolipid species at (G) 6 h and (H) 12 h.

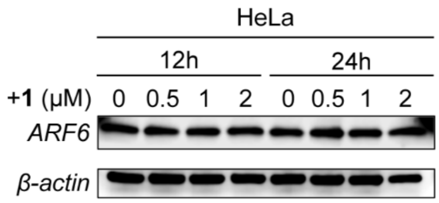


**Fig. S30. Total ARF6 protein levels are unchanged with whorl accumulation.** Immunoblotting of ARF6 in HeLa cells treated with vehicle or **1**.

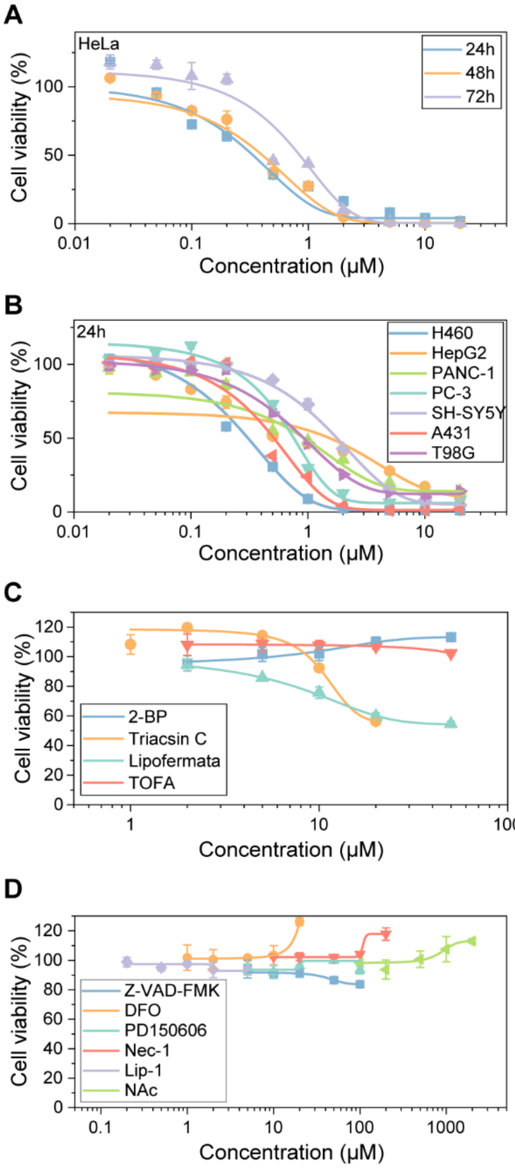


**Fig. S31. Cell viability of cells treated with 1 or lipid-pathway inhibitors alone.**
(A) Cell viability of HeLa cells treated with **1**. (B) cell viability of other cancer cell lines treated with **1**. (C, D) Cell viability of HeLa cells treated with the indicated inhibitors alone, for 24 h.

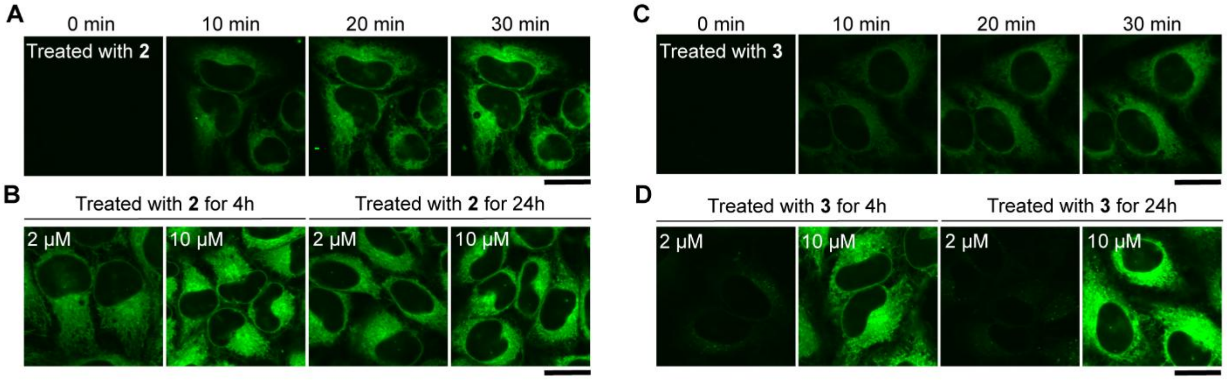


**Fig. S32. Peptide analogs lacking the thioester warhead (2, 3) do not induce whorls.**
(A, C) Time-lapse and (B, D) CLSM images of HeLa cells treated with compound **2** (A, B) or compound **3** (C, D). (Scale bar = 20 μm)

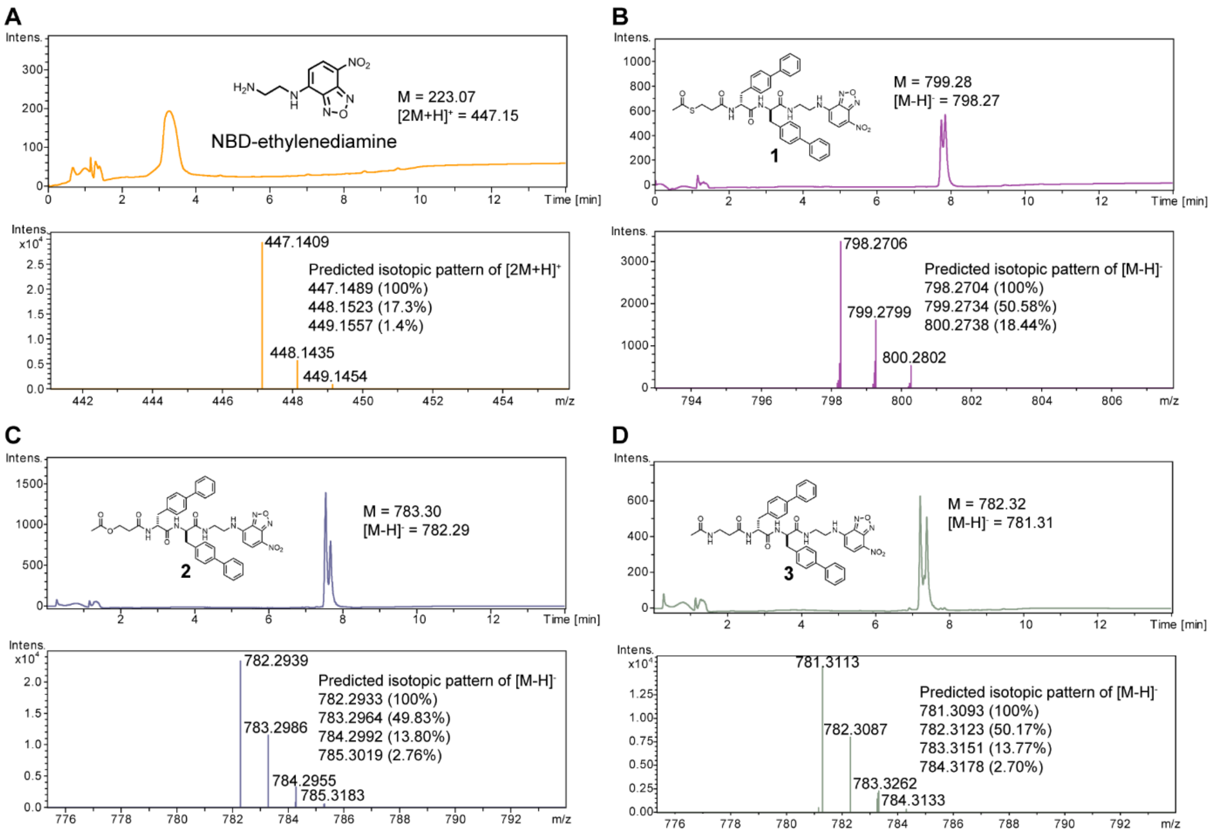


**Fig. S33. High resolution mass spectrum of compounds.**
High resolution mass spectrum of (A) NBD-ethylenediamine, (B) AcS-bb-NBD (**1**), (C) AcO-bb-NBD (**2**) and (D) AcN-bb-NBD (**3**).

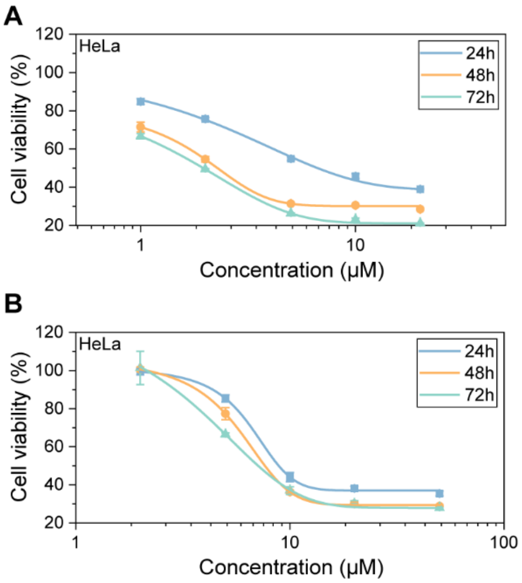


**Fig. S34. Cell viability of HeLa cells treated with compounds 2 and 3.**
(A) Cell viability of HeLa cells treated with compound **2**. (B) Cell viability of HeLa cells treated with compound **3**.

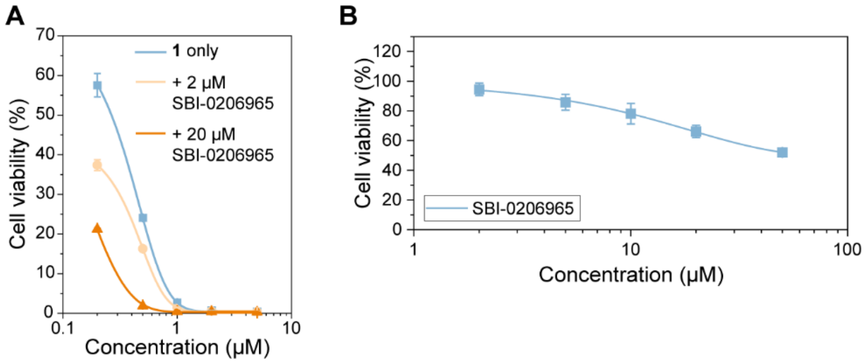


**Fig. S35. ULK1 inhibition with SBI-0206965 enhances the cytotoxicity of 1.** Cell viability of HeLa cells treated with SBI-0206965 for 24 h, alone or with **1**.

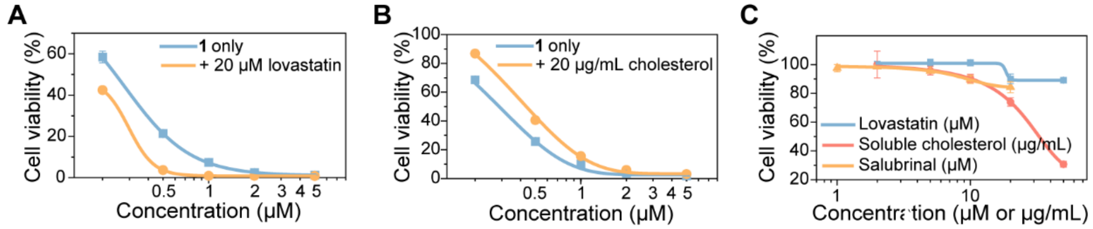


**Fig. S36. Cell viability of HeLa cells treated with lovastatin, cholesterol, or salubrinal alone.** (A, B) Cell viability of HeLa cells treated with **1** in the presence or absence of lovastatin or water-soluble cholesterol. (C) Cell viability of HeLa cells treated with lovastatin or water-soluble cholesterol alone for 24 h.

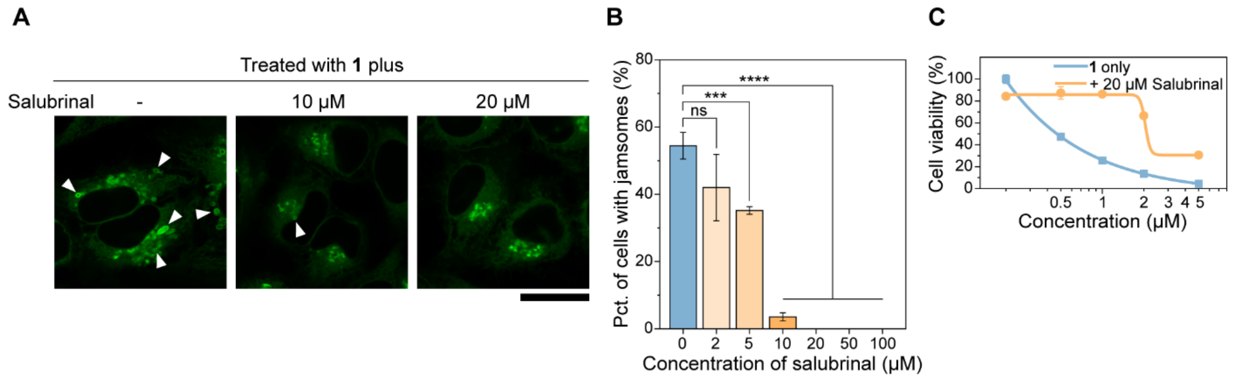


**Fig. S37. Salubrinal reduces whorl abundance and the toxicity of 1.**
(A) CLSM images and (B) quantification of whorls in cells pretreated with salubrinal for 30 min, followed by co-treatment of **1** and salubrinal for 4 h. (C) Cell viability of HeLa cells treated with **1** in the presence or absence of salubrinal for 24 h. (Scale bar = 20 μm) White arrowheads indicate whorls.

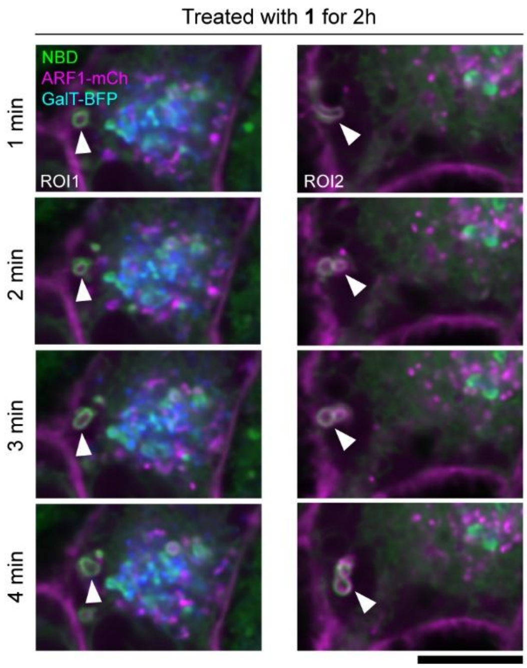


**Fig. S38. Whorls form near the plasma membrane.**
Time-lapse CLSM images of HeLa cells co-transfected with ARF1-mCherry and GalT-BFP and treated with **1** for 2 h, showing whorls near the plasma membrane. (Scale bar = 10 μm) White arrowheads indicate whorls.
